# An efficient early-pooling protocol for environmental DNA metabarcoding

**DOI:** 10.1101/2022.02.15.480497

**Authors:** Masayuki Ushio, Saori Furukawa, Hiroaki Murakami, Reiji Masuda, Atsushi J. Nagano

## Abstract

Environmental DNA (eDNA) metabarcoding, a method that applies high-throughput sequencing and universal primer sets to eDNA analysis, has been a promising approach for efficient, comprehensive biodiversity monitoring. However, significant money-, labor-, and time-costs are still required for performing eDNA metabarcoding. In the present study, we assessed the performance of an “early-pooling” protocol (a protocol based on the 1st PCR indexing) to reduce the experimental costs of the library preparation for eDNA metabarcoding. Specifically, we performed three experiments to test the effects of 1st PCR and 2nd PCR indexing protocols on the community composition revealed by eDNA metabarcoding, of post-1st-PCR exonuclease purification on index-hopping, and of the number of PCR replicates and eDNA template volume on the number of detected OTUs. By analyzing 204 eDNA libraries from 3 natural aquatic ecosystems and 1 mock eDNA sample, we show that (i) the 1st PCR indexing does not cause clear biases in the outcomes of eDNA metabarcoding, (ii) post-1st-PCR exonuclease purification reduces the risk of index-hopping, and (iii) increasing the eDNA template volume can increase the number of detected OTUs and reduce the variations in detected community compositions, as can increasing the number of the 1st PCR replicates. Our results show that an early-pooling protocol with post-1st-PCR exonuclease purification and an increased amount of DNA template will reduce the risk of index-hopping, the costs for consumables and reagents, and the handling time in the library preparation, and that it produces comparable results to a 2nd-PCR-indexing protocol. Therefore, once a target metabarcoding region is determined and a set of indexed-1st-PCR primers is prepared, the early-pooling protocol provides a cost-, labor-, and time-efficient way to process a large number of samples.

## Introduction

Frequent and comprehensive ecosystem monitoring is a basis for effective biodiversity conservation, resource management and near future forecasting. Direct visual census, direct capture (e.g., fishing and insect collection), and camera/video trapping (e.g., for forest mammals) have traditionally been used as tools for biodiversity monitoring (Masuda *et al*. 2010; Samejima *et al*. 2012; Nakagawa 2019). These data are invaluable and contribute to better understanding, conservation, and management of ecosystems under intense human pressures. However, these methods are usually time-consuming and require professional expertise such as taxonomic identification skill, which prevents their application to a frequent, large spatial-scale biodiversity monitoring.

Environmental DNA (eDNA), namely, DNA isolated from environmental samples without capturing target organisms, has been used to detect the presence of macro-organisms (Taberlet *et al*. 2012; e.g., Miya *et al*. 2015; Yamamoto *et al*. 2017; Taberlet *et al*. 2018). In the case of macro-organisms, eDNA originates from various sources such as metabolic waste or damaged tissue (Kelly *et al*. 2014), and the eDNA contains information about the species identity of organisms that produced it. Since the first application of eDNA analysis to natural ecosystems (Ficetola *et al*. 2008), eDNA has been used in many studies as a tool for investigation of the distributions of fish species in ponds, rivers and seawater (Jerde *et al*. 2011; Minamoto *et al*. 2011; Sigsgaard *et al*. 2015; Ushio *et al*. 2018), as well as the distributions of other aquatic/semiaquatic/terrestrial organisms (Deiner *et al*. 2016; Bista *et al*. 2017; Ishige *et al*.2017;Ushio *et al*. 2017; Yonezawa *et al*. 2020).

eDNA metabarcoding, a method that applies universal primer sets and high-throughput sequencing to eDNA analyses, has now been widely used for comprehensive sequencing of target metabarcoding regions in an eDNA sample (Taberlet *et al*. 2012; Kelly *et al*. 2014; Miya *et al*. 2015; Ushio *et al*. 2017; Yamamoto *et al*. 2017). For example, a previous study demonstrated that an eDNA metabarcoding using fish-targeting universal primers (MiFish primers) enabled detection of more than 230 fish species from seawater in a single study (Miya *et al*. 2015). Another study demonstrated that eDNA metabarcoding can detect nearly 300 families from river eDNA samples and that the families contain both aquatic and terrestrial organisms, which suggested that a river collects and transports eDNA from surrounding area and that river eDNA can provide a large amount of information about the biodiversity in the watershed (Deiner *et al*. 2016).

Although eDNA metabarcoding is a powerful and promising method for comprehensive and efficient biodiversity monitoring, it has significant costs. While sequencing costs have been continuously declining over the past few decades (for example, see https://www.genome.gov/about-genomics/fact-sheets/DNA-Sequencing-Costs-Data), sample collection, DNA extraction and library preparation are still laborious and time-consuming. To reduce the costs of eDNA metabarcoding, researchers have been developing technologies that enable automated sampling and rapid DNA extraction (Yamahara *et al*. 2019; Fukuzawa *et al*. 2021), but a commonly used 2-step PCR protocol for eDNA library preparation remains laborious (e.g., Minamoto *et al*. 2021).

The 2-step PCR protocol includes the first round PCR (1st PCR) that amplifies a target metabarcoding region using a universal primer (e.g., mitochondrial 12S region by MiFish primers), followed by the second-round PCR (2nd PCR) that appends sample-specific index sequences to the amplicons (i.e., the 2nd PCR indexing). After the 2nd PCR, multiple samples may be combined (i.e., multiplexing, or “pooling”) because samples can be distinguished by the sample-specific index sequences afterward. However, until the index sequences are appended, each sample should be separately processed, and careful operations are required to prevent cross-contaminations. Alternatively, a sample-specific index may be appended at the 1st PCR (i.e., the 1st PCR indexing), an approach which has been used in metabarcoding studies targeting prokaryotes (Caporaso *et al*. 2011; Smets *et al*. 2016) and eukaryotes (Leray & Knowlton 2017; Zizka *et al*. 2019; Minardi *et al*. 2022). This protocol enables multiplexing after the 1st PCR (i.e., “early-pooling”). However, such an approach could introduce index-specific amplification biases (Berry *et al*. 2011; O’Donnell *et al*. 2016), leading eDNA researchers to prefer the 2nd PCR indexing protocol despite its relatively high experimental costs. In addition, the early-pooling might increase the risk of index hopping (Esling *et al*. 2015) if the samples are pooled without inactivating the primers and enzyme in the reaction.

Furthermore, because the concentration of target eDNA is often low, especially for macro-organisms (e.g., fish), researchers have recommended preparing multiple technical replicates at the 1st PCR to maximize the species detection probabilities (e.g., >4-8 replicates: Doi *et al*. 2019), which inevitably increases the handling time and costs for plastic consumables and reagents. Increasing the number of the 1st PCR replicates, however, increases the total volume of template eDNA used in the PCR. For example, 8 replicates of a 1st PCR which uses 1 *μ*l of template eDNA per replicate uses 8 *μ*l of template eDNA in total. Therefore, although increasing the number of the 1st PCR replicates is currently recommended, it is still unclear whether the thus-far observed patterns that the species detection probability increased with the number of replicates employed in the 1st PCR is due to the increased number of replicates or the increased volume of template DNA.

Because techniques in molecular biology have been continuously improving, whether common recommendations still hold for the current situation and how outcomes change using updated reagents and protocols are often unclear. For such reasons, in the present study, we assessed the performance of an “early-pooling” protocol (i.e., a protocol based on the 1st PCR indexing) to reduce the experimental costs of the library preparation for eDNA metabarcoding. Using indexed 1st PCR primers, updated reagents, and several customized protocols, we performed fish eDNA metabarcoding for natural and mock samples and compared the results with those revealed by a common 2nd-PCR-indexing protocol. Specifically, we performed three experiments to test the effects of (i) indexing methods on the community composition revealed by eDNA metabarcoding (Experiment I), (ii) post-PCR exonuclease purification on index-hopping (Experiment II), and (iii) the number of PCR replicates and volumes of eDNA template on the number of detected OTUs (Experiment III). Based on the experiments, we discuss the advantages and disadvantage of the early-pooling protocol and propose an efficient protocol for eDNA metabarcoding library preparation which is applicable to a large number of eDNA samples.

## Mateirals and Methods

### Safeguarding against potential contaminations during the sampling and experiments

Prior to the DNA extractions and library preparation, work-space and equipment were sterilized. Filtered pipet tips were always used to safeguard against cross-contamination.

### Preparation of eDNA samples and standard fish DNAs

Water samples for the method testing were collected from 3 natural environments: 2 marine and 1 freshwater environments in Japan. Seawater samples were collected at Nagahama, Maizuru in Kyoto prefecture (35°29′24″ N, 135°22′6″ E) on 13 May 2021 and Otomi, Takahama in Fukui prefecture (35°32′24″ N, 135°30′3″ E) on 14 May 2021. Seawater was collected by throwing a bucket tied with a rope from a pier 10 times. Three Sterivex filter cartridges (SVHV010RS, Merck Millipore, Darmstadt, Germany) were used at each site, and 1,000 ml of seawater was filtered by using a disposal syringe for each cartridge, followed by the addition of RNAlater (ThermoFisher Scientific, Waltham, Massachusetts, USA) to prevent DNA degradation. The cartridges were transferred to the laboratory within 30 min and stored at −20°C.

Freshwater samples were collected from the Seta River in Otsu, Shiga prefecture, Japan (34°57′39″ N, 135°54′32″ E) on 30 March 2020. We collected 1,600 ml of river water using plastic bottles, and filtered the water sample using eight Sterivex filter cartridges (each Sterivex filtered 200 ml of water sample). The cartridges were brought back to the laboratory within 1 hour, and stored at −20°C. The samples were kept at 4°C during transport.

Detailed protocols for the DNA extractions are described in the Supplementary Methods. Briefly, DNA was extracted using a DNeasy Blood & Tissue kit (Qiagen, Hilden, Germany) following a protocol described previously (Miya *et al*. 2016). After the cell lysis and purification, DNA was eluted using 100 *μ*l of the elution buffer. Extracted DNAs from the same study site were combined and treated as one composite sample for the method testing. Eluted DNA samples were stored at −20°C until further processing.

In addition to the 3 samples from natural ecosystems, we prepared 10 fish-like standard DNAs. These 10 fish-like standard DNAs were synthesized using gBlocks Gene Fragments service by Integrated DNA Technologies, Inc. (Coralville, IA, USA). Equal amounts of the 10 standard DNAs were mixed as Standard DNA Mix (STD_Mix) and treated identically with the eDNA samples from the 3 study sites. More information about the standard fish DNAs is available in the Supplementary Methods and Table S1.

### Quantification and normalization of fish eDNA concentrations

Before we started the experiments, the concentrations of fish eDNA and the standard DNAs were normalized so that similar numbers of sequence reads could be generated for each site and each replicate. Fish eDNA concentrations were estimated using quantitative PCR (qPCR) as described in the Supplementary Methods. The minimum concentration of fish eDNA was found in the river eDNA sample (43.2 copies/*μ*l), and thus the other eDNA samples were diluted so that their fish eDNA concentrations are 43.2 copies/*μ*l.

### Experiment I: Effects of indexing methods on the community composition revealed by eDNA metabarcoding

#### Two indexing methods and two DNA polymerases

In Experiment I, a primary objective was to test the effects of 2 indexing methods, the 1st PCR indexing and 2nd PCR indexing, on the detected fish eDNA compositions. The 1st PCR indexing is a method to append a sample-specific index sequence in the 1st PCR (Fig. 1a). Similar approaches were previously applied in eukaryotic eDNA metabarcoding studies (O’Donnell *et al*. 2016; Leray & Knowlton 2017; Zizka *et al*. 2019; Minardi *et al*. 2022). A 1st PCR primer set for the 1st PCR indexing (hereafter, “the 1st-indexing-1st-PCR primer”) is composed of Illumina sequencing primer, 8-base sample-specific index, and MiFish-U primer (Table 1). In the 1st PCR indexing protocol, the PCR products are pooled and purified after the 1st PCR, and the Illumina P5/P7 adapters are appended for the pooled sample at the 2nd PCR using “the 1st-indexing-2nd-PCR primer” (Table 1). On the other hand, the 2nd PCR indexing protocol does not append a sample-specific index at the 1st PCR and uses “the 2nd-indexing-1st-PCR primer” to amplify a target region (Fig. 1a: e.g., Miya *et al*. 2015). After the 1st PCR, each PCR product is separately purified, and is then used as a template for the 2nd PCR. At the 2nd PCR, sample-specific index sequences and the Illumina P5/P7 adapters are appended to the amplicon.

**Figure 1.**
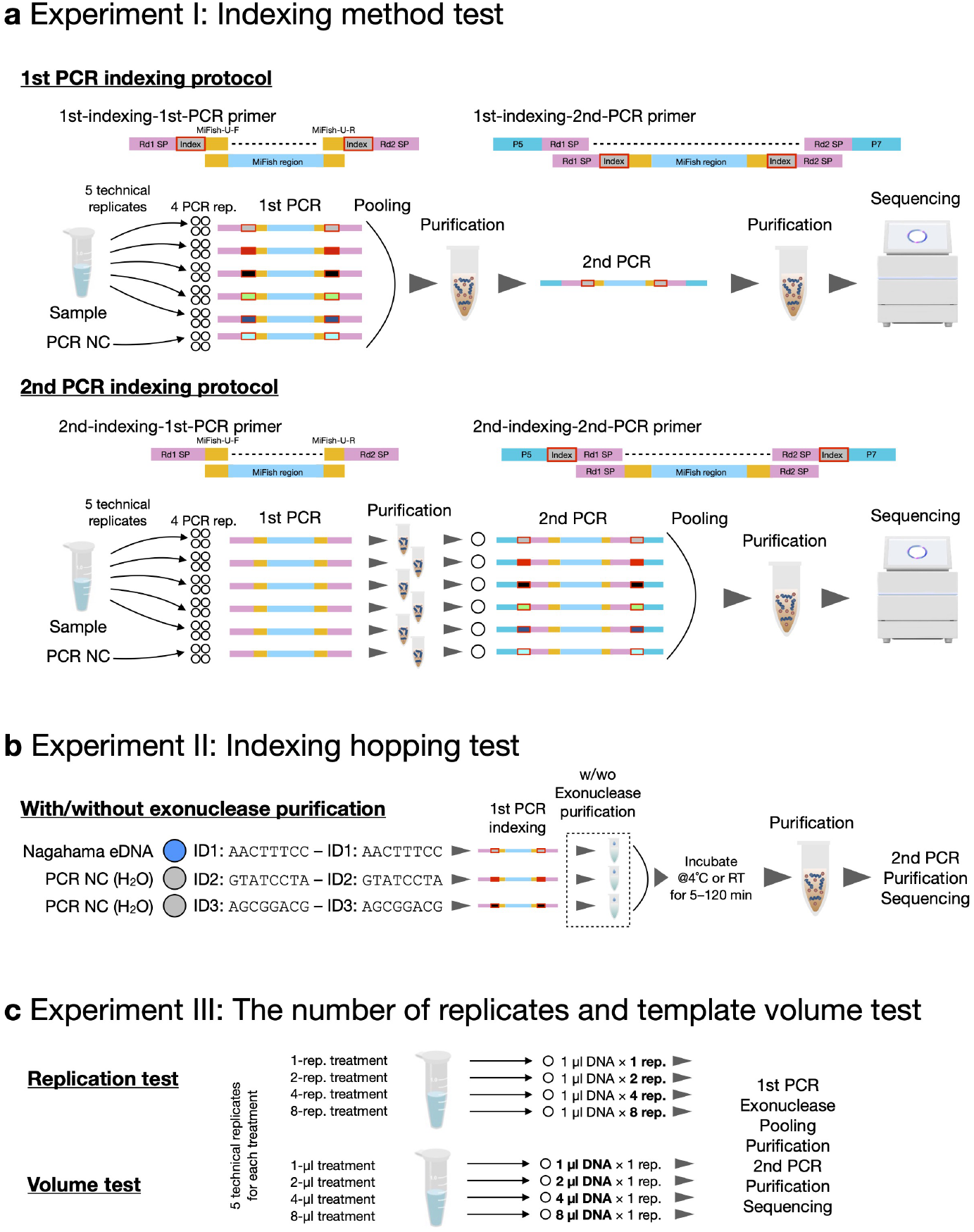
Experimental designs of this study. (**a**) Experiment I. Effects of the indexing method were tested. A 1st PCR indexing protocol appends sample-specific index sequences in the 1st PCR using “1st-indexing-1st-PCR primers” while a 2nd PCR indexing protocol appends the index sequences in the 2nd PCR using “2nd-indexing-2nd PCR primers.”“Purification” indicates magnetic beads purification in Experiment I. (**b**) Experiment II. Effects of exonuclease purification on index-hopping events were tested. Exonuclease purification was performed for each sample. Then samples were combined and incubated to test the effects of incubation time and temperature. (**c**) Experiment III. The effects of the 1st PCR replicates and template DNA volume were tested. Other processes were identical with that of Experiment II with exonuclease purification.

**Table 1.**
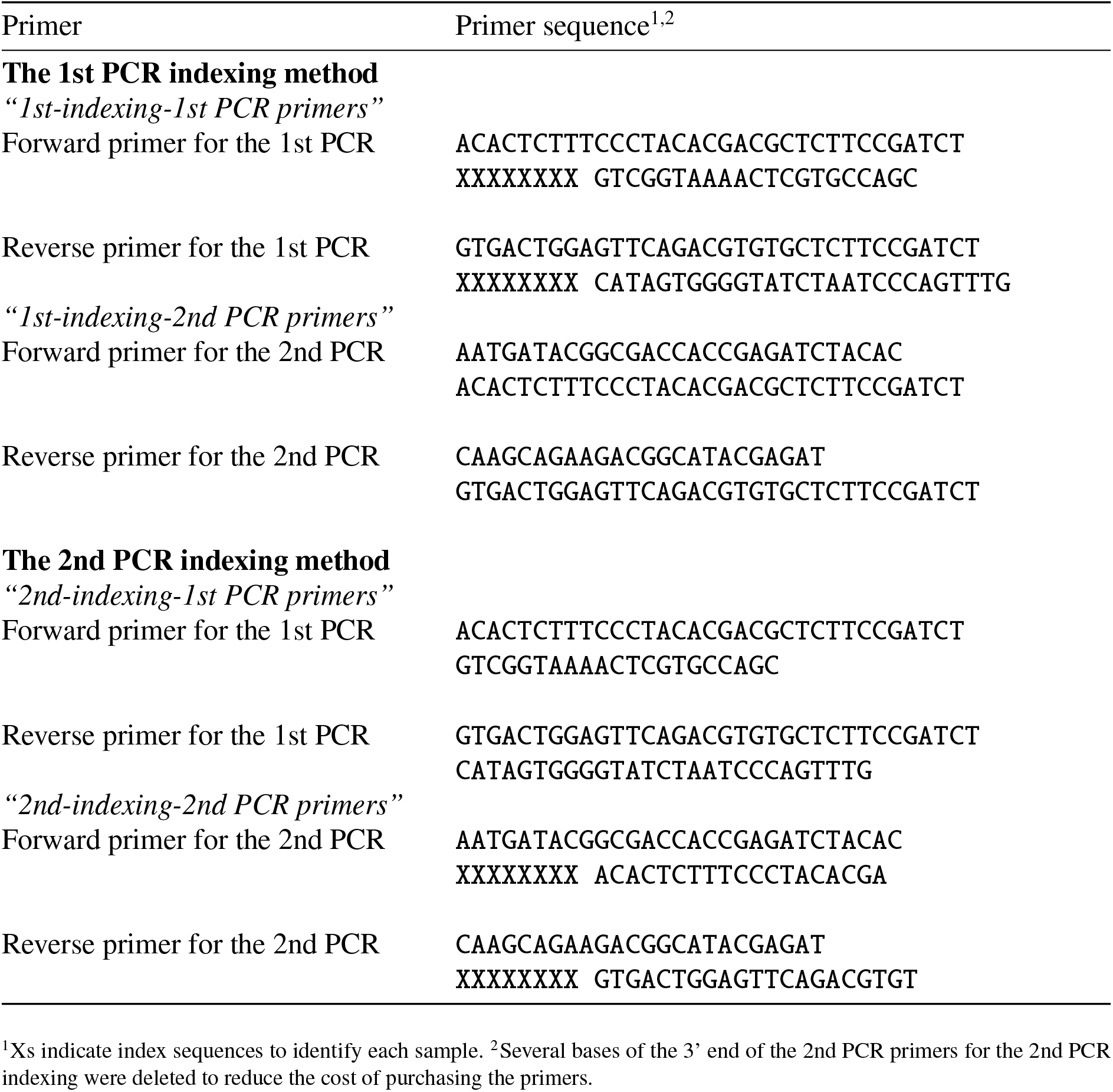
Primer sequences used in the present study.

As a secondary objective, we evaluated effects of two DNA polymerases on the outcomes of eDNA metabarcoding: KAPA HiFi HotStart ReadyMix (KAPA Biosystems, Wilimington, WA, USA; hereafter, “KAPA”) and Platinum SuperFi II PCR Master Mix (ThermoFisher Scientific, Waltham, MA, USA; hereafter, “Platinum”). The former is a commonly used enzyme in eDNA metabarcoding, while the latter is a newer enzyme with higher fidelity according to the manufacturer’s information.

#### Experimental design

In Experiment I, we tested 3 library preparation protocols: The 2nd PCR indexing with KAPA, the 2nd PCR indexing with Platinum, and the 1st PCR indexing with Platinum. The first two were for comparing effects of enzymes, and the last two were for comparing effects of indexing methods. In each library preparation protocol, eDNA samples originated from 3 study sites (i.e., Nagahama, Otomi, and Seta-river) and one standard DNA mix were analyzed. In each study site, 5 technical replicates and 1 negative control (H_2_O) were included. Thus, we had 72 eDNA libraries in total, that is, (5 technical replicates +1 negative control) × (3 study sites and 1 standard DNA) × (3 protocols) =72.

#### Library preparations and iSeq sequencing

Thermal cycle profiles of the 1st and 2nd PCR are described in Table S2. We used dual-unique combinations of forward and reverse indices for different templates (samples) for Illumina sequencing, which reduces the risk of index-hopping (Esling *et al*. 2015). All combinations of indices are available in https://github.com/ong8181/eDNA-early-pooling/tree/main/sampledata.

For all treatments, we performed the 1st and 2nd PCR following the manufactures’ protocols as described in the Supplementary Methods. Briefly, for “the 2nd PCR indexing with KAPA” treatment (i.e., a common 2-step PCR protocol), KAPA HiFi HotStart ReadyMix was used for the 1st and 2nd PCR. Sample-specific index sequences were appended in the 2nd PCR. For “the 2nd PCR indexing with Platinum” treatment, Platinum SuperFi II PCR Master Mix was used instead of KAPA, and sample-specific index sequences were appended in the 2nd PCR. For “the 1st PCR indexing with Platinum” treatment, Platinum was used instead of KAPA, and sample-specific index sequences were appended in the 1st PCR. After the 2nd PCR, the libraries were purified, target-sized DNA was excised, the double-stranded DNA concentrations of the libraries were quantified, and the 3 libraries were combined as one sample. The double-stranded DNA concentration of the combined library was then adjusted to 50 pM using 10 mM Tris-HCl (pH 8.5) and the DNA was then sequenced by the iSeq 100 system (Illumina, San Diego, CA, USA) using iSeq 100 Reagent v2 (2 × 150 bp PE). For the sequencing, 30% PhiX was spiked-in to improve the sequencing quality.

### Experiment II: Effects of post-1st-PCR purification on index-hopping events

The early-pooling protocol could cause index-hopping events because the enzyme and indexed-primers were not inactivated before the sample pooling. Therefore, in Experiment II, we explicitly measured the frequency of index-hopping events in the 1st PCR indexing treatment and evaluated how the exonuclease purification, temperature, and time after the sample pooling affect the frequency of index-hopping events (Fig. 1b).

#### Experimental design

In Experiment II, the frequency of index-hopping events was evaluated by preparing 1 positive sample and 2 negative samples for each treatment. eDNA extracted from Nagahama seawater was used as the positive sample, while H_2_O was used as the negative samples. These 3 samples were distinguished by dual-unique index, for example, the combinations of ID1–ID1, ID2–ID2, and ID3–ID3 (Fig. 1b). In the 1st PCR indexing protocol in Experiment I, 1st PCR products were immediately combined after the 1st PCR and the polymerase and primers were not inactivated when the samples were pooled. Therefore, we may find “unused” index combinations such as ID1–ID2, ID2–ID1, ID1–ID3, and ID3–ID1, which is a signature of index-hopping events. Two-time index-hopping may generate the combinations of ID2–ID2 and ID3–ID3, but we disregard the probability because it should be much lower than that of one-time index hopping.

We evaluated effects of 3 variables on the frequency of index-hopping events: exonuclease purification for each 1st PCR product (hereafter, “with exonuclease” treatment) and incubation temperature and time after the sample pooling. The purification for each 1st PCR product was performed by incubating the PCR products using ExoSAP-IT Express (ThermoFisher Scientific, Waltham, MA, USA), which is less labor-intensive than purification using magnetic beads. The 3 1st PCR products for each treatment were then pooled and incubated at 2 temperatures (on ice [ca. 4°C] or at room temperature [ca. 22°C]) for 3 durations (5 min, 30 min, and 120 min). Thus, we had 36 eDNA libraries in total, that is, (1 positive sample +2 negative controls) × (with/without exonuclease) × (2 temperatures) × (3 durations) =36.

#### Library preparations and iSeq sequencing

Detailed protocols for Experiment II are described in the Supplementary Methods. For 36 samples, the 1st PCR was performed by the 1st PCR indexing protocol using Platinum as in Experiment I with some modifications. Briefly, for the “with exonuclease” treatment only, 4 *μ*l of ExoSAP-IT Express was added to each 10-*μ*l PCR product. The mixture was incubated for 4 min at 37°C, followed by the incubation for 1 min at 80°C. In addition, we performed the 2nd PCR using “2nd-indexing-2nd-PCR primers” to append additional indices to the library (i.e., quad-index approach). This approach was adopted because the 3 indices used to distinguish samples in each treatment were identical among the treatments.

### Experiment III: Effects of the number of PCR replicates and volume of eDNA template on the number of OTUs detected

As the Results of Experiments I and II suggested that fish community compositions revealed by the 1st PCR indexing with the exonuclease purification were comparable to those revealed by the 2nd PCR indexing protocol, in Experiment III, we evaluated effects of the replications and template volume in the 1st PCR to further reduce the potential experimental- and labor-cost of the library preparation.

#### Experimental design

In Experiment III, we prepared 8 treatments, that is 1, 2, 4, and 8 replicate treatments (“replicate” treatment) and 1, 2, 4, and 8-*μ*l treatment (“volume” treatment), for 2 sample types (Nagahama seawater and standard DNA samples) (Fig. 1c). In each treatment, 5 technical replicates and 1 negative control were included. We chose the Nagahama sample because its eDNA diversity was highest among the 4 sample types (ca. 20-30 fish species were detected in Experiment I and II). A standard DNA sample was included because we know the number of DNA species in it *a priori* (i.e., 10 species should be detected). We had 96 eDNA libraries in total, that is, (5 technical replicates +1 negative control) × (4 replicate treatments and 4 volume treatments) × (2 sample types) =96.

#### Library preparations and iSeq sequencing

Detailed protocols are described in the Supplementary Methods. DNA libraries were prepared using the 1st PCR indexing protocol using Platinum as in Experiment I with the following modifications. Briefly, for all treatments, exonuclease purification for each 1st PCR product was included. For “replicate” treatment, the number of 1st PCR replicates was changed for each treatment. For “volume” treatment, the total reaction volume was increased from 10 *μ*l to 20 *μ*l so that we could include up to 8 *μ*l of eDNA template. As in Experiment II, “2nd-indexing-2nd-PCR primers” were used to append additional indices to the library (i.e., quad-index approach). This approach was adopted to distinguish libraries with identical 1st PCR indices.

### Sequence data processing

Detailed procedures for our sequence data processing are described in the Supplementary Methods. Our sequence data includes sample-specific indices inside the sequencing primers and we prepared a custom shell script using seqkit 2.1.0 (Shen *et al*. 2016) to demultiplex our samples (see https://github.com/ong8181/eDNA-early-pooling/tree/main/01_Demultiplex and https://doi.org/10.5281/zenodo.6045851). Raw sequences were demultiplexed and MiFish primer regions were trimmed by cutadapt 2.10 (Martin, 2011). The quality of our sequence data was high, and after primer trimming, 8,948,113 sequence reads (%> Q30 =93.9; average 43,863 reads per sample) remained for the three experiments.

The demultiplexed, primer-trimmed sequences were processed using DADA2 (Callahan *et al*. 2016), an amplicon sequence variant (ASV) approach, for each iSeq run. First, at the quality filtering process, low quality and unexpectedly short reads were removed. Error rates were learned, and sequences were dereplicated, error-corrected, and merged to produce an ASV-sample matrix. Then, chimeric sequences were removed. ASVs detected in the three experiments were merged and clustered into OTU at 97% similarity using DECIPHER package of R (Wright 2016), which converted the ASV-sample matrix into the OTU-sample matrix. Taxonomic identification was performed for OTUs based on the query-centric auto-k-nearest-neighbor (QCauto) method (Tanabe & Toju 2013) implemented in Claident v0.9.2021.10.22 (https://www.claident.org/). Because the QCauto method requires at least 2 sequences from a single taxon, standard DNAs were separately identified using BLAST (Camacho *et al*. 2009).

### Statistical analyses and data visualization

All statistical analyses and visualizations were performed using a free statistical environment, R 4.1.2 (Team 2021). The sample metadata, taxa information assigned to each OTU, and OTU-sample matrix were imported as an object using phyloseq package of R (McMurdie & Holmes 2013). The merged phyloseq object was then divided into the three experiments and analyzed separately. In the present study, we describe the results of statistical tests using the term “statistical clarity” rather than “statistical significance” to avoid misinterpretations, according to the recommendations by Dushoff *et al*. (2019).

#### Statistical analyses of Experiment I

For Experiment I, the data set was first divided into each sample type (i.e., Nagahama, Otomi, Seta-river, or standard DNA; Fig. S1) after a brief inspection of the sequence reads distribution. Then, the sequence reads were rarefied by the minimum number of sequence reads detected in each sample type to remove the effects of sequence depth on results. Rarefying sequence reads by the minimum number of sequence reads is usually not recommended for ecological studies (McMurdie & Holmes 2014). However, in our case this approach was suitable because replicates in each sample type were technical replicates, and because we wanted to focus on the effects of the library preparation protocols on the outcomes by removing the effects of sequence depth. This type of rarefaction was also applied in Experiment III.

After the rarefaction within each sample type, the fish community compositions were first visualized using barplots. The effects of the library preparation protocols on the numbers of detected OTUs were tested using the generalized linear model (GLM assuming Poisson distribution). The dependence of fish community compositions on the library preparation protocols was assessed by nonmetric dimensional scaling (NMDS) based on the Bray-Curtis dissimilarity, and the differences in the detected community compositions among the library preparation protocols were tested by the analysis of similarities (ANOSIM) using vegan package of R (Oksanen *et al*. 2008). Also, the effects of the library preparation protocols on the Bray-Curtis dissimilarities were tested by bootstrap analysis (see https://github.com/ong8181/eDNA-early-pooling/tree/main/08_Exp1_1st2nd for detail).

#### Statistical analyses of Experiment II

For Experiment II, there were 1 positive sample and 2 negative samples in each treatment. Because each treatment has 3 dual-unique indices (ID1, ID2, and ID3), there are 9 possible combinations of indices (3 × 3 combinations), among which 4 “unused” combinations indicate possible index-hopping events (i.e., ID1-ID2, ID2-ID1, ID1-ID3, and ID3-ID1). Sequence reads assigned to OTUs with these 4 indices were standardized by the sequence reads of the positive sample. For example, in the case that OTU001 has 10,000 reads in the positive sample (ID1-ID1) while 10 reads are detected as index-hopped (e.g., ID1-ID2 combination), the proportion of index-hopped reads is calculated as 0.1%. The pattern was visualized and the statistical clarity of 3 variables, i.e., purification process, temperature, and time, were tested by GLM.

#### Statistical analyses of Experiment III

For Experiment III, the data set was first divided into each sample type (i.e., Nagahama, or standard DNA). Then, the sequence reads were rarefied to the minimum number of sequence reads in each sample type to remove the effects of sequence depth. Then, the effects of the reaction scale (i.e., the number of 1st PCR replicates and the volume of template DNA) were statistically tested using GLM. In addition, how the number of 1st PCR replicates and template DNA volume influence the detection of rare OTUs was evaluated (see Results). The effects of the number of replicates and total template volume on the Bray-Curtis dissimilarities were tested by bootstrap analysis as described for Experiment I.

#### Additional statistical analyses

We performed additional analyses to investigate the index- and the protocol-specific biases in the detected community composition using data from Experiment I, II, and III. For the index-specific bias test, samples sequenced using the identical index were grouped, and the variations in the relative abundance of dominant OTUs among the index sequences were tested. For the protocol-specific bias test, we visualized how the protocols changed the fish eDNA compositions of the Nagahama samples because the Nagahama samples were analyzed using various library preparation protocols (e.g., the 1st PCR indexing method, the 2nd PCR indexing method, and with/without exonuclease purification).

### Code and Data availability

Sequence data are deposited in DDBJ Sequence Read Archives (DRA) (DRA accession number =DRA013399). Analysis scripts and the information about R packages are available at Github (https://github.com/ong8181/eDNA-early-pooling/).

## Results

In total, 8,948,113 sequence reads were generated from 204 samples, of which 8,530,488 reads remained after the DADA2 sequence processing. Of the sequences that remained, 8,518,552 reads were assigned to 113 fish OTUs, including the standard DNAs. A small number of sequence reads (11,936 reads; 0.13%) was assigned to 21 non-fish OTUs such as mouse (*Mus musculus*), pig (*Sus scrofa*), and common pochard (*Aythya ferina*). In the subsequent analysis, we focused on the 113 fish OTUs.

### Experiment I: Effects of indexing methods on the community composition revealed by eDNA metabarcoding

In total, 1,910,880 reads were generated from 72 samples in Experiment I. To archive fair comparisons between the protocols, we rarefied sequence reads to the minimum number of sequence reads within each sample type (21,452 reads for Nagahama; 8,351 reads for Otomi; 10,151 reads for Seta river; 9,574 reads for standard DNA mix). The sequence reads generated in the negative controls were minor compared with those generated in the samples (Fig. S1).

Overall, we detected very similar OTU richness and community compositions regardless of the library preparation protocols (Figs. 2 and S2). For example, the effect of the protocols on the fish OTU richness was statistically unclear (Fig. S2a; *P* > 0.05). *Acanthopagrus* sp (OTU002), *Hexagrammos* sp. (OTU092), and Cyprinidae (OTU004) were the most dominant fish OTUs in Nagahama, Otomi, and Seta river, respectively, and similar community compositions were detected by the 3 protocols (Fig. S2b). Similarly, the NMDS plot shows that the samples are largely overlapped among the protocols (Fig. 2a-d). There was no clear statistical difference in the fish community composition between “the 2nd PCR indexing with Platinum” and “the 2nd PCR indexing with KAPA” (ANOSIM; *P* > 0.05). Similarly, there was no clear statistical difference between “the 1st PCR indexing with Platinum” and “the 2nd PCR indexing with Platinum” (ANOSIM; *P* > 0.05). We further investigated the reproducibility in each protocol (i.e., variations in the technical replicates within a protocol) in the fish community compositions by calculating Bray-Curtis dissimilarity. We found that the 1st PCR indexing increased the dissimilarities among technical replicates in the standard DNA samples (Fig. 2e; *P* < 0.001). On the other hand, for the other samples, differences in the dissimilarities among technical replicates were statistically unclear (Fig. 2e; *P* > 0.05). In addition, the relative abundance of the dominant fish OTUs were not different among the protocols (Fig. S3; *P* > 0.05) except that of the second most dominant OTU (*Parablennis*) in the Nagahama samples.

**Figure 2.**
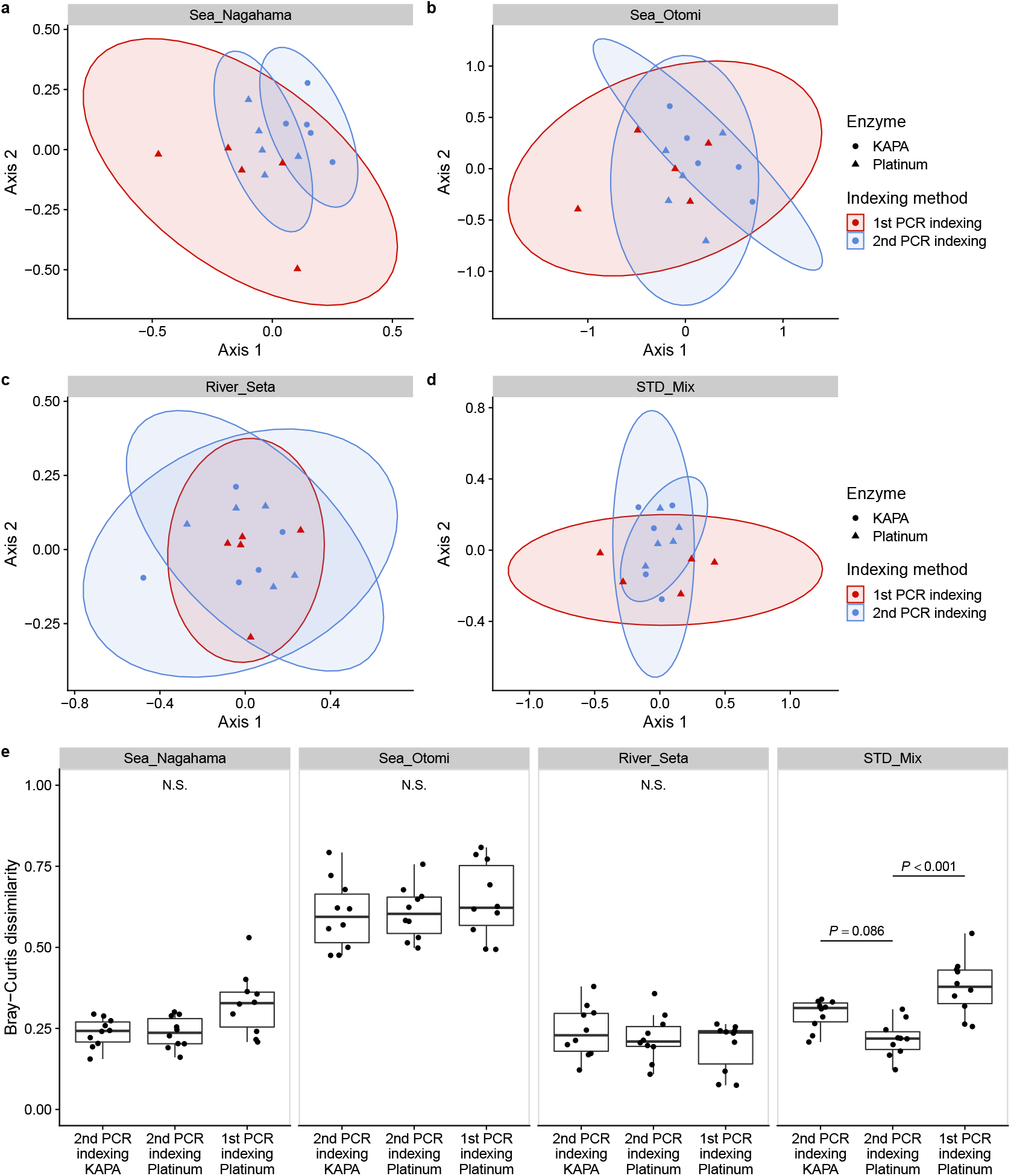
Nonmetric dimensional scaling (NMDS) and Bray-Curtis dissimilarities of fish eDNA composition detected in Experiment I. Fish eDNA compositions of (**a**) seawater samples collected in Nagahama, Kyoto, Japan, (**b**) seawater samples collected in Otoumi, Fukui, Japan, (**c**) freshwater samples collected in Seta River, Otsu, Japan, and (**d**) a mixture of 10 standard fish DNAs. Filled circles and triangles indicate that KAPA HiFi HotStart ReadyMix and Platinum SuperFi II PCR Master Mix was used for PCR, respectively. Red and blue circles indicate that sample-specific index sequences were appended by the 1st PCR indexing method and the 2nd PCR indexing method, respectively. (**e**) Each panel represents eDNA samples collected in one study site. Points in each panel represent Bray-Curtis dissimilarities of fish eDNA compositions between two eDNA samples. The thick bar indicates the median value of the Bray-Curtis dissimilarities in each treatment. Statistical clarity was tested by bootstrap test (see Methods).

### Experiment II: Effects of post-PCR clean-up on index-hopping

In total, 3,060,823 reads were generated from 36 samples in Experiment II. We found relatively frequent index-hopping events for the non-purified treatments (Fig. 3). Although the proportion of sequence reads for each OTU generated by index-hopping is generally low (i.e., about 0.01-0.3%), “unused” combinations of index sequences were found in many OTUs in the without-exonuclease treatments (red and dark red points in Fig. 3). On the other hand, the frequency of index-hopping events was dramatically reduced when the 1st PCR products were purified using exonuclease before sample pooling (blue and light blue points in Fig. 3). No index-hopping events were observed with the exonuclease treatment within 5 min after sample pooling at 4°C and room temperature. Some index-hopping events were observed with the exonuclease treatment 30 and 120 min after sample pooling at 4°C and room temperature, but the number of OTUs of which indices were hopped was much lower than that with non-purified treatments. The effect of exonuclease was statistically clear (binominal GLM; *P* < 0.0001), while those of time and temperature were not (P >0.05).

**Figure 3.**
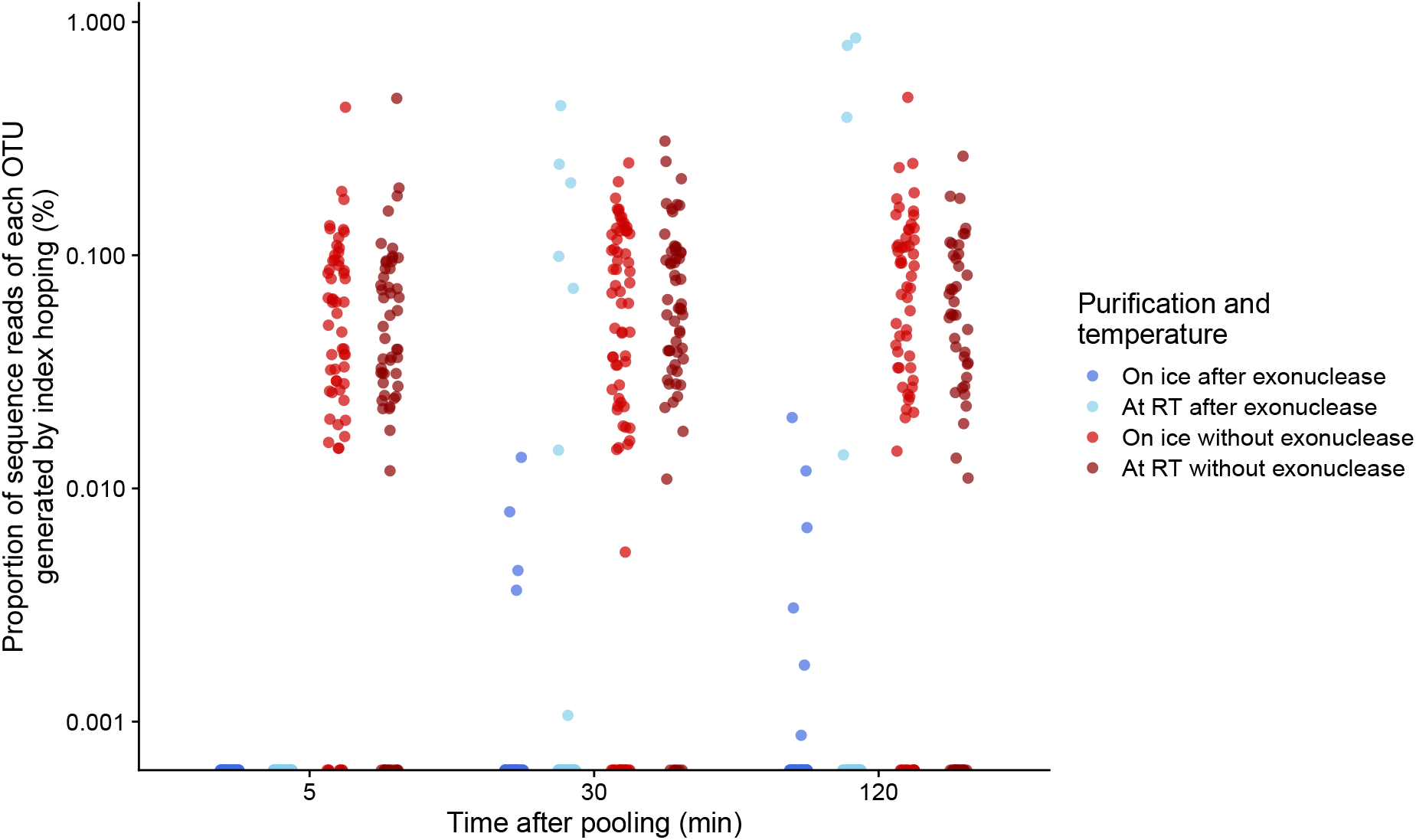
The estimation of the index-hopping probability of each library preparation method. *y*-axis indicates the proportion of sequence reads for each OTU generated by index-hopping. We used three index combinations in Experiment II, i.e., ID1_F (AACTTTCC)-ID1_R (AACTTTCC) (positive sample), ID2_F (GTATCCTA)-ID2_R (GTATCCTA) (H_2_O), and ID3_F (AGCGGACG)-ID3_R (AGCGGACG) (H_2_O). Index-hopping is defined as the occurrence of any of the four “unused” index combinations in this study: ID1_F-ID2_R, ID1_F-ID3_R, ID2_F-ID1_R, and ID3_F-ID1_R. We ignored the other possible combinations as described in the main text. Each point represent the index-hopped sequence reads for each OTU divided by the sequence reads of the corresponding OTU detected in the positive sample. Blue and light blue points indicate that exonuclease-purified 1st PCR products were pooled and incubated on ice and at room temperature (22°C), respectively. Red and dark red points indicate not-purified 1st PCR products were pooled and incubated on ice and at room temperature (22°C), respectively.

### Experiment III: Effects of the number of 1st PCR replicates and volume of eDNA template on the number of OTUs detected

In total, 3,546,849 reads were generated from 96 samples in Experiment III, and we rarefied sequence reads to the minimum number of sequence reads within the sample type (15,394 reads for Nagahama; 3,025 reads for standard DNA mix). One replicate in the 1-rep. treatment was excluded from the analysis because only 304 reads were detected from the replicate. Overall, the community compositions detected in the treatments were similar (Fig. S4), and the numbers of OTUs in Nagahama seawater and standard DNA samples were increased with the number of 1st PCR replicates and volumes of eDNA template (Fig. 4a, b).

**Figure 4.**
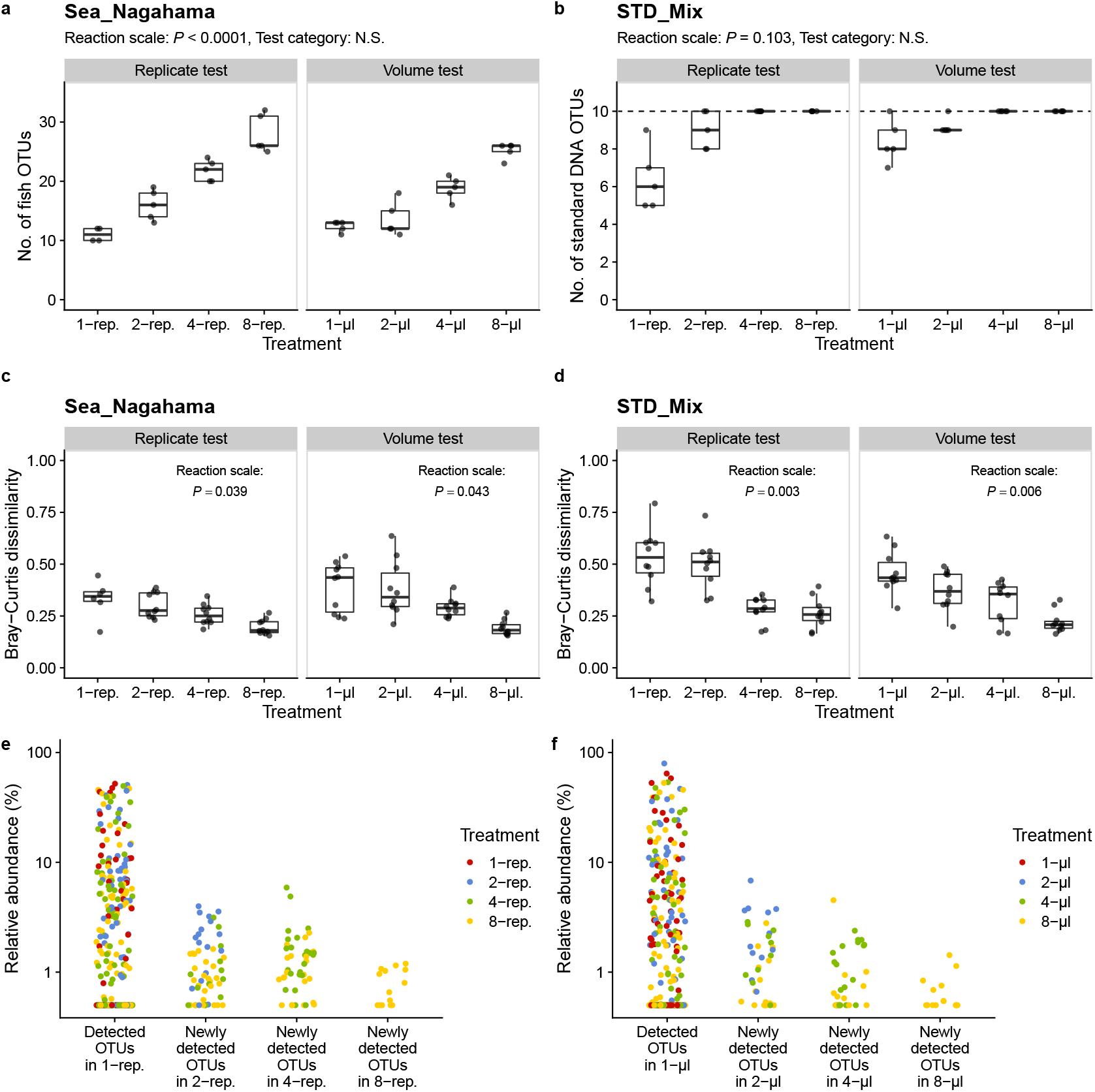
Effects of the number of replicates and the volume of template DNA in the 1st PCR reaction on community compositions and OTU richness. (**a**, **b**) Effects of the number of replicates and the template DNA volume on the number of fish OTUs in the Sea_Nagahama samples (**a**) and standard DNA samples (**b**). In **b**, Dashed horizontal line indicates the number of standard DNAs included in the reactions. Statistical clarity was tested by GLM. (**c**, **d**) Effects of the number of replicates and the template DNA volume on Bray-Curtis dissimilarities in the Sea_Nagahama samples (**c**) and standard DNA samples (**d**). Statistical clarity was tested by the bootstrap analysis. (**e**, **f**) The relationship between the relative abundance of OTUs and detected treatments. In e, each point indicates that each OTU is newly detected in the replication treatment described on x-axis. Points at “Detected OTUs in 1-rep.” indicate all OTUs detected in the 1-rep. treatment. Colors indicate the treatment in which OTUs are detected. (**f**) Results for the volume test of the Sea_Nagahama samples.

For Nagahama seawater samples, in the replicate test, 11.0 ±1.2 (mean ±standard deviation), 16.0 ±2.6, 21.8 ±1.8, and 28.0 ±3.2 fish OTUs were detected in 1-rep., 2-rep., 4-rep., and 8-rep. treatments, respectively (Fig. 4a). In the volume test, 12.4 ±0.9, 13.6 ±2.9, 18.8 ±1.9, and 25.2 ±1.3 fish OTUs were detected for Nagahama samples in 1-*μ*l, 2-*μ*l, 4-*μ*l, and 8-*μ*l treatments, respectively (Fig. 4a). For Nagahama samples, the effect of reaction scale on the number of fish OTUs was statistically clear (Poisson GLM; *P* < 0.0001) while that of test category (“replicate” or “volume”) was not.

For standard DNA samples, in the replicate test, 6.4 ±1.7 and 9.0 ±1.0 fish OTUs were detected in 1-rep. and 2-rep. treatments, respectively (Fig. 4b). In the volume test, 8.4 ±1.1 and 9.2 ±0.4 fish OTUs were detected in 1-*μ*l and 2-*μ*l treatments, respectively (Fig. 4b). For the other treatments, all 10 standard fish DNAs were detected in all replicates. For standard DNA samples, the effect of reaction scale was statistically unclear (Poisson GLM; *P* =0.103) possibly because of the small number of fish standard DNAs included in the samples.

As for the reproducibility of each treatment (i.e., within-treatment variations in the detected community compositions), increasing the number of the 1st PCR replicates or template volumes statistically clearly reduced the Bray-Curtis dissimilarities (Fig. 4c, d; *P* < 0.05). For both Nagahama and standard DNA samples, 8-rep. and 8-*μ*l treatments showed more consistent community compositions among the technical replicates than 1-rep. and 1-*μ*l treatments did (Fig. 4c, d).

We further investigated how the number of 1st PCR replicates and the template DNA volumes influence the OTU detection using Nagahama samples (standard DNA samples showed similar results, but are not shown because they have low fish OTU numbers). We did this by identifying OTUs that were newly detected in a more-replicated treatment or a larger template-volume treatment (Fig. 4e, f). For example, “Detected OTUs in 2-rep” indicates that the OTUs were not detected in the 1-rep. treatment, but were newly detected in the 2-rep. treatment (Fig. 4e). Newly detected OTUs in more-replicated treatments generally had lower abundance than OTUs detected in 1-rep. treatment (Fig. 4e), which indicates that rare OTUs can be detected more easily by increasing the number of 1st PCR replicates. These patterns were almost the same for the template volume test (Fig. 4f). Newly detected OTUs in larger template-volume treatments generally had lower abundance than OTUs detected in 1-*μ*l treatment (Fig. 4f).

### Testing index- and protocol-specific biases

Lastly, using the data from Experiment I, II, and III, we tested index- and protocol-specific biases in the eDNA metabarcoding. Index-specific biases were tested by quantifying the relative abundance of dominant OTUs detected in the Nagahama seawater samples (Fig. 5a). OTU002, OTU007, and OTU056 were the 3 most abundant taxa, and were assigned as *Acanthopagrus* sp. (most likely, Japanese black seabream, *A. schlegelii*), *Parablennius yatabei* (Yatabe blenny), and *Takifugu* sp. (most likely, pufferfish, *T. alboplumbeus*), respectively, which are all common fish species in the region (Masuda 2008; Masuda *et al*. 2010). The relative abundances of the 3 OTUs estimated by a common method, the 2nd-indexing protocol, were 39.5%, 18.4%, and 6%, respectively (red dashed horizontal lines in Fig. 5a). Technical replicates were grouped into each index sequence in Experiment I, II, and III, ignoring detailed experimental steps in each Experiment (e.g., with/without exonuclease, the number of 1st PCR replicates, and the volume of template DNA). We found similar relative abundances of the top 3 OTUs with that of the common 2nd-indexing protocol regardless of the index sequences, and the effect of index sequence on the relative abundance of the 3 OTUs was statistically unclear (binomial GLM; *P* > 0.05; tested only for Experiment III).

**Figure 5.**
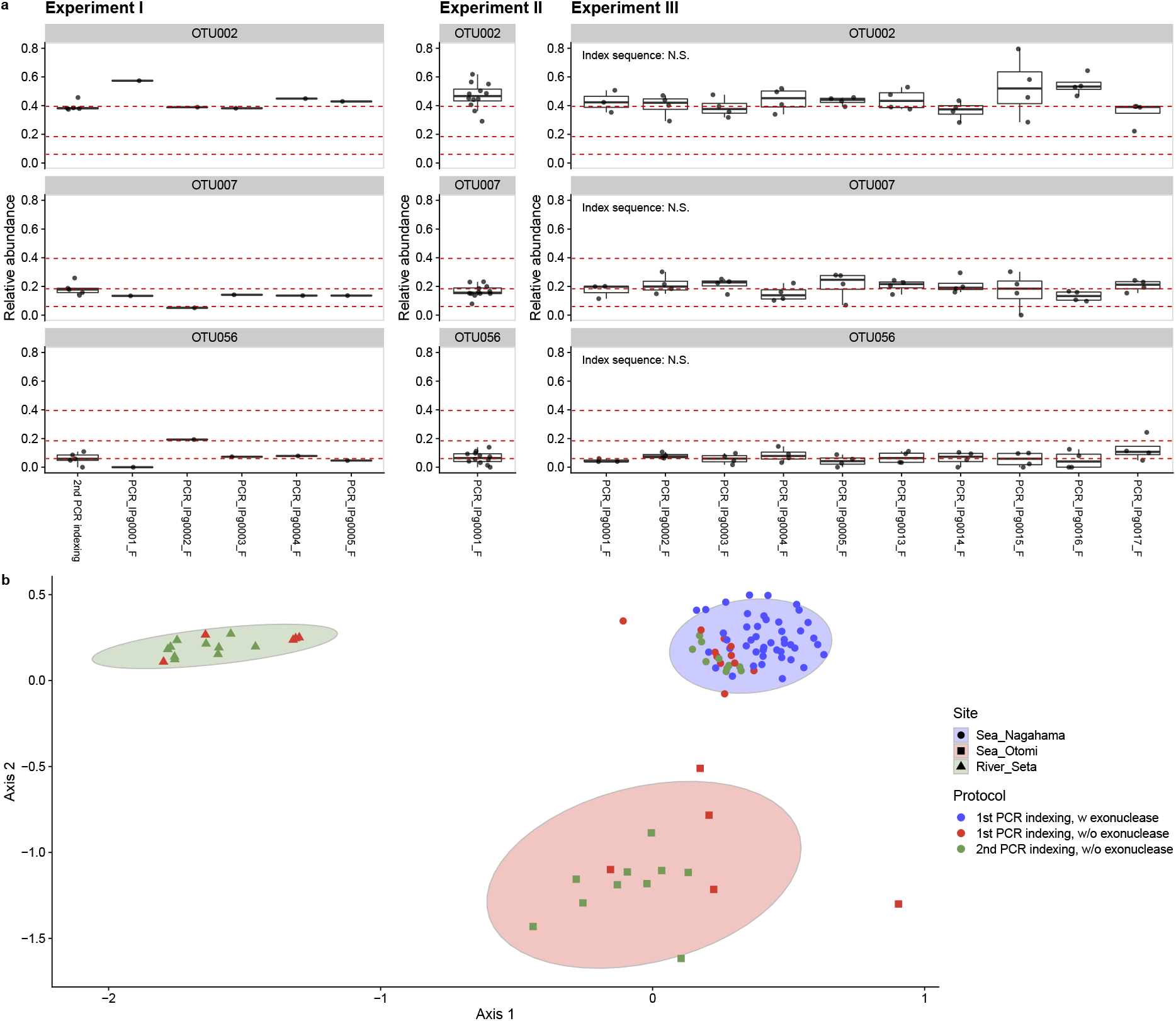
Effects of index sequences and protocols on the relative abundance and community compositions of fish detected in the Nagahama samples. (**a**) *y*-axis shows the relative abundance of three most-dominant OTUs. *x*-axis shows library preparation method (= “2nd PCR indexing”) or the names of index sequence used in the 1st PCR. Red dashed horizontal lines indicate the mean relative abundance of each OTU detected by the 2nd PCR indexing method. In Experiment I, only one library was sequenced using each index. In Experiments II and III, multiple libraries were sequenced using the same index. Note that different experimental treatments (e.g., incubation time and temperature in Experiment II) are grouped in this figure. In Experiment III, statistical clarity was tested by GLM. (**b**) The effects of library preparation protocols on the fish community compositions of three study sites visualized by Nonmetric dimensional scaling (NMDS). All Nagahama samples are clearly distinguished from natural eDNA samples from the other study sites. Symbols and colors indicate the library preparation protocols and study sites, respectively. Ellipses indicate 95% confidential intervals for each study site. An NMDS plot with more detailed sample information is shown in Figure S5b.

Protocol-specific community compositions were tested by leveraging the results from Ex-periment I, II, and III. Fish community compositions of the Nagahama seawater samples were grouped into each protocol and visualized in Figs. 5b and S5. In general, the fish community compositions were similar regardless of the protocols used (Fig. S5a). The community compositions seemed to be variable in less-replicated treatments (1-rep. and 2-rep. treatment) or smaller template-volume treatments (1-*μ*l and 2-*μ*l treatments), but the variations became smaller in more-replicated or larger template-volume treatments (Fig. S5a; also evident in Fig. 4c, d). The patterns were also evident in NMDS (Figs. 5b and S5b). Some experimental steps (e.g., exonuclease purification after the 1st PCR) could affect the community composition slightly, but the fish community compositions could be clearly distinguished from those detected in different sample types, such as river and other seawater samples (Figs. 5b and S5b).

## Discussion

### Effects of the indexing method on the fish community compositions

In Experiment I, we showed that the 1st PCR indexing and the early-pooling did not cause significant protocol-specific biases in the community compositions or the relative abundance of dominant OTUs (Figs. 2, 5, and S1-S3), which is contrary to the findings of previous studies (Berry *et al*. 2011; O’Donnell *et al*. 2016). O’Donnell *et al*. (2016) employed shorter indexed-primers (31 bases including 3 ambiguous bases and 6 unique index bases) for the 1st PCR indexing than we did (62-69 bases including 8 unique index bases; Table 1), which causes a larger variation in *T_m_* among the primers. As differences in *T_m_* induce primer-template binding efficiency (Wu *et al*. 2009), this difference might be a reason for the discrepancy. Several previous studies also reported little differences in the outcome between the 2 indexing protocols (Leray & Knowlton 2017; Zizka *et al*. 2019), which is consistent with our study. Although it is difficult to elucidate the details of the mechanisms that generated the differences between the studies, the 1st PCR indexing generates qualitatively and quantitatively similar results to those generated by the 2nd PCR indexing, at least in our protocol.

### Effects of post-1st-PCR purification on the index-hopping events

In Experiment II, we found that post-1st-PCR exonuclease purification reduced the risk of cross-contaminations (Fig. 3). It is known that index-hopping is one event that prevents accurate (e)DNA-based evaluations of ecological community compositions (Esling *et al*. 2015), and how to reduce the risk of index-hopping events is an important issue. This is particularly important in an early-pooling protocol because primers and enzymes are still “active” at the time of pooling. In our experiments, index-hopping events were evident as detections of “unused” combinations of index sequences (Fig. 3), but the post-1st-PCR exonuclease purification reduced the risk. Although adding the purification step increases the cost and handling time for the library preparation, the exonuclease-based purification is much easier and more rapid compared with magnetic beads-based purification. Also, it does not require an expensive instrument (e.g., a microplate washer such as Hydrospeed, TECAN) even for a large number of samples. Furthermore, the pooling steps can be performed at room temperature, as this temperature did not increase the risk of the index-hopping events,.

### Effects of the number of 1st PCR replicates and template volume on OTU richness

In Experiment III, we showed that increasing the number of the 1st PCR replicates and template DNA volume had similar effects on the detected OTU richness (Fig. 4a, b). Our analysis found that rarer OTUs tended to be detected more easily in the more-replicated and larger template-volume treatments (Fig. 4e, f), suggesting that the positive effects of increasing the number of the 1st PCR replicates and template DNA volume are simply due to the increased probability of the occurrence of rare OTUs in the reaction. In addition, increasing the number of the 1st PCR replicates and template DNA reduces variations among technical replicates (Fig. 4c, d), which could be due to the mitigation of random sampling effects of rare DNAs. In a previous study, increasing the number of the 1st PCR replicates was recommended to increase the eDNA-based detection probability of fish species (Doi *et al*. 2019). However, this inevitably increases the number of pipettings and the amount of consumables used in the experiment (e.g., pipette tips and PCR tubes), which will increase the cost of consumables and reagents and the handling time of the library preparations. On the other hand, increasing template volume does not increase the cost or handling time, and thus would be preferable to increasing the number of replicates, especially when a large number of samples are processed.

### A recommended protocol

We showed that (i) the 1st PCR indexing does not cause clear biases on the outcomes, at least in our experimental settings (i.e., with our reagents and primer structure), (ii) post-1st-PCR exonuclease purification reduces the risk of the index-hopping events, and (iii) increasing the template DNA volume can increase the OTU richness detected and reduce variations in detected community composition among technical replicates, as increasing the number of the 1st PCR replicates does. Based on these results, we recommend the following protocol for the eDNA metabarcoding library preparation:

1. Prepare indexed-1st PCR primers (e.g., 48 or 96 indexed primers).
2. Append index sequences at the 1st PCR (the 1st-PCR indexing). At the 1st-PCR, the number of the 1st PCR replicate(s) may be one if the template DNA volume is 4-8 *μ*l or greater.
3. Perform post-1st-PCR purification by exonuclease to reduce the risk of index-hopping.
4. Pool the purified 1st PCR products.
5. If necessary, adjust DNA concentration using magnetic beads.
6. Perform the 2nd PCR to append sequencing adapter for one composite sample (the 2nd indices may be appended to increase the maximum number of multiplexed samples; i.e., “quad-index approach”).
7. Purify the PCR products, adjust concentrations, and sequence DNA.

This “early-pooling” protocol reduces the costs for consumables, reagents, and handling time (see Table S3 for the cost and time estimations). In addition, the “quad-index approach” enables multiplexing a large number of samples easily. For example, the combination of 96 1st-PCR indices and 96 2nd-PCR indices enables distinguishing nearly 10,000 samples (96 × 96 = 9216) at a single sequencing run, which would contribute to large-scale eDNA monitoring.

### Potential limitations of the early-pooling protocol

Despite several advantages of the early-pooling eDNA metabarcoding protocol, there are several potential limitations/disadvantages. First, the early-pooling protocol requires indexed 1st-PCR primers for each metabarcoding region, which will potentially increase the money cost for an indexed primer set. Thus, researchers who try to test many different metabarcoding regions for a relatively small number of samples may first consider using the common 2nd-PCR indexing protocol which uses the same indexed primer set for any metabarcoding region. Second, increasing the template DNA volume will simultaneously increase the concentrations of PCR inhibitors such as humic substances (Schrader *et al*. 2012). For samples with a high concentration of PCR inhibitor, one may consider increasing the number of the 1st PCR replicates or using a PCR-inhibitor-tolerant DNA polymerase.

## Conclusions

Environmental DNA metabarcoding is an emerging tool for efficient biodiversity monitoring, yet sample processing, that is, sample collection, DNA extraction, and library preparation for sequencing, still requires costs and effort. In the present study, we have shown that an early-pooling protocol with post-1st-PCR exonuclease purification and an increased amount of DNA template will reduce the risk of index-hopping and costs for consumables, reagents and handling time in the library preparation, and that it produces comparable results with a common 2nd-PCR-indexing protocol. Therefore, once a target metabarcoding region is determined and a set of indexed-1st-PCR primers is prepared, the early-pooling protocol provides a cost-, labor-, and time-efficient way to process a large number of samples, which will contribute to understanding and near-future forecasting of ecosystem dynamics.

## Endnotes

### Data and code availability

All scripts used in the present study are available in Github (https://github.com/ong8181/eDNA-early-pooling). Sequence data are deposited in DDBJ Sequence Read Archives (DRA) (DRA accession number =DRA013399).

## Acknowledgements

We thank Misaki Shiomi in Maizuru Fisheries Research Station for assisting with water collection and eDNA extraction. This research was supported by KAKENHI (B) 20H03323 (to MU) and the Hakubi Project in Kyoto University (to MU).

## Conflict of Interest Statements

S.F. is employed by Clockmics, which is undertaking eDNA library preparations and sequencing. The commercial affiliation of the author (S.F.) does not alter our adherence to the journal policies on sharing data and materials. None of the authors will benefit directly financially from the publication of this paper.

## Author Contributions

MU and AJN conceived and designed research; MU, SF, HM, and RM collected test samples; SF and MU performed experiments; MU analyzed data; MU, SF, and AJN wrote the first draft; All authors discussed the results and completed the manuscript.

