## Supplementary Methods for "An efficient early-pooling protocol for environmental DNA metabarcoding"

### Contents

- Supplementary Methods

### Supplementary Methods

#### DNA extractions

DNA was extracted using a DNeasy Blood & Tissue kit (Qiagen, Hilden, Germany) following a protocol described in a previous study (Miya *et al.* 2016) with some modifications.

For Nagahama and Otomi seawater samples, RNAlater solution (ThermoFisher Scientific, Waltham, MA) was removed from the outlet of the filter cartridge by centrifuging it at 4000 g for 2 min. One ml of $H_2O$ was added to wash the filter cartridge, followed by centrifugation at 4000 g for 2 min to remove water. Then, Proteinase K solution (20 $\mu$l) and buffer AL (200 $\mu$l) were mixed, and the mixture was added to each filter cartridge which contained ca. 200 $\mu$l of $H_2O$ ($H_2O$ that remained after the centrifugation). The materials on the cartridge filters were subjected to cell lysis by incubating the filters on a rotary shaker (20 rpm; Roller 6 digital, IKA-Werke Staufen, Germany) at 56°C for 20 min. The incubated and lysed mixture was transferred into a new 3-ml tube from the inlet (not the outlet) of the filter cartridge by centrifugation (4,000 g for 2 min). The collected DNA was purified using a DNeasy Blood & Tissue kit following the manufacturer's protocol. After the purification, DNA was eluted using 200 $\mu$l of the supplied elution buffer. Extracted DNAs from the same study site were combined and treated as one composite sample for the method testing. Eluted DNA samples were stored at -20°C until further processing.

For Seta-River samples, first, Proteinase K solution (20 $\mu$l), PBS (220 $\mu$l) and buffer AL (200 $\mu$l) were mixed, and 440 $\mu$l of the mixture was added to each filter cartridge. The materials on the cartridge filters were subjected to cell lysis by incubating the filters on a rotary shaker (15 rpm; DNA oven HI380R, Kurabo, Osaka, Japan) at 56°C for 10 min. The incubated and lysed mixture was transferred into a new 2-ml tube from the inlet (not the outlet) of the filter cartridge by centrifugation (3,500 g for 1 min). The collected DNA was purified using a DNeasy Blood & Tissue kit following the manufacturer's protocol. After the purification, DNA was eluted using 100 $\mu$l of the supplied elution buffer. Extracted DNAs from multiple filter cartridges were combined and treated as one composite sample for the method testing. Eluted DNA samples were stored at -20°C until further processing.

#### Preparation of standard fish DNAs

Ten fish-like standard DNAs were synthesized using gBlocks Gene Fragments service by Integrated DNA Technologies, Inc. (Coralville, IA, USA). The ten standard sequences were designed based on fish mitochondrial 12S metabarcoding region (MiFish region: Miya *et al.* 2015). Five of them were designed so that they have no mismatch with fish-targeted universal primers, MiFish-U-F/R (Miya *et al.* 2015), and their lengths are 220 bases including the primer regions (insert length = 172 bases). Also, they have the identical conserved regions with real fish species, but randomized sequences in variable regions. The other five standard DNAs were designed so that they are similar to specific fish genera (i.e., *Anguilla*, *Cyprinus*, *Engraulis*, *Lateolabrax*, and *Takifugu*), and their lengths were 217–221 bases including the primer region (insert length = 169–173 bases). They have no or a few mismatches in the forward primer region while they have no mismatch in the reverse primer region. Similar to the first five standard DNAs, they have the identical conserved regions with real fish species, but randomized sequences in variable regions. The standard DNA sequences are provided in Table S1.

Overall, the 10 standard fish DNAs have similar or identical primer and conserved regions with existing fish species, so we expected that their PCR amplification efficiencies would be similar to those of real fish DNAs. Sequences in the variable regions of the standard DNAs were randomized, and as a result, the standard DNAs are at most *ca.* 80% similar to the MiFish region of existing fish (the expected differences in sequences between the standard DNAs and real fish DNAs are 34-35 bases), which makes it easy to distinguish the standard DNAs from real fish DNAs. These features are useful for testing cross-contaminations between samples. Also, the standard DNAs were used as positive samples in the present study (i.e., we should detect 10 species if a metabarcoding method correctly detects eDNA diversity). Equal amounts of the 10 standard DNAs were mixed as Standard DNA Mix (STD_Mix) and treated identically with the eDNA samples from the three study sites.

**Quantifications and normalizations of fish DNA concentrations**

Before we started the experiments, the concentrations of fish eDNA and the standard DNA were normalized so that similar sequence reads could be generated for each site and each replicate. Fish eDNA concentrations were estimated using quantitative PCR (qPCR) using Light Cycler 480 System II (Roche, Basel, Switzerland). Briefly, qPCR of the MiFish region was performed using MiFish primer set (Miya *et al.* 2015) (without Illumina sequencing primer, adapter and sample-specific index sequences). One $\mu$l of each DNA was added to a qPCR reaction containing 4 $\mu$l of 2.5 $\mu$M forward primer, 4 $\mu$l of 2.5 $\mu$M reverse primer, 10 $\mu$l of Platinum SuperFi II PCR Master Mix (ThermoFisher Scientific, Waltham, MA, USA), and 1 $\mu$l of 20 × EvaGreen (Biotium, San Francisco, CA, USA). The thermal cycle profile after an initial 30 s denaturation at 98°C was as follows (60 cycles): denaturation at 98°C for 10 s; annealing at 60°C for 10 s; and extension at 72°C for 15 s (fluorescent measured at this step). The minimum concentration of fish eDNA was found in the river eDNA sample (43.2 copies/$\mu$l), and thus the other eDNA samples were diluted so that their fish eDNA concentrations are 43.2 copies/$\mu$l.

**Experiment I: Library preparation protocols and iSeq sequencing**

For "the 2nd PCR indexing with KAPA" treatment, the 1st PCR was carried out with a 10-$\mu$l reaction volume containing 5.0 $\mu$l of 2 × KAPA HiFi HotStart ReadyMix, 1.2 $\mu$l of each 2.5 $\mu$M 2nd-indexing-1st-PCR F/R primer (final concentration was 0.3 $\mu$M each as recommended by the manufacturer), 1.6 $\mu$l of sterilized distilled $H_2O$, and 1.0 $\mu$l of template. We performed four 1st PCR replicates, and those replicates were pooled to mitigate the PCR dropouts. The pooled four 1st-PCR products for each sample were separately purified using AMPure XP (PCR product: AMPure XP beads = 1:0.8; Beckman Coulter, Brea, CA, USA). The purified and 10-fold diluted 1st PCR products were used as templates for the 2nd PCR. The 2nd PCR was carried out with a 20-$\mu$l reaction volume containing 10 $\mu$l of 2 × KAPA HiFi HotStart ReadyMix, 2.4 $\mu$l of each 2.5 $\mu$M 2nd-indexing-2nd-PCR F/R primer (final concentration was 0.3 $\mu$M each), 3.2 $\mu$l of sterilized distilled $H_2O$ and 2.0 $\mu$l of diluted 1st PCR product. After the 2nd PCR, the indexed PCR products were combined.

For "the 2nd PCR indexing with Platinum" treatment, the 1st PCR was carried out with a 10-$\mu$l reaction volume containing 5.0 $\mu$l of 2 × Platinum SuperFi II PCR Master Mix, 2.0 $\mu$l of each 2.5 $\mu$M 2nd-indexing-1st-PCR F/R primer (final concentration was 0.5 $\mu$M each as recommended by the manufacturer), and 1.0 $\mu$l of template. Four 1st PCR replicates were pooled to mitigate the PCR dropouts. The pooled four 1st-PCR products for each sample were separately purified using AMPure XP (PCR product: AMPure XP beads = 1:0.8). The purified and 10-fold diluted 1st PCR products were used as templates for the 2nd PCR. The 2nd PCR was carried out with a 20-$\mu$l reaction volume containing 10 $\mu$l of 2 × Platinum SuperFi II PCR Master Mix, 4.0 $\mu$l of each 2.5 $\mu$M 2nd-indexing-2nd-PCR F/R primer (final concentration was 0.5 $\mu$M each), and 2.0 $\mu$l of diluted 1st PCR product. After the 2nd PCR, the indexed PCR products were combined.

For the “1st PCR indexing with Platinum” treatment, the 1st PCR was carried out with a 10-$\mu$l reaction volume containing 5.0 $\mu$l of 2 × Platinum SuperFi II PCR Master Mix, 2.0 $\mu$l of each 2.5 $\mu$M 1st-indexing-1st-PCR F/R primer, and 1.0 $\mu$l of template. Four 1st PCR replicates were pooled to mitigate the PCR dropouts. The pooled four 1st-PCR products were then combined as one sample, which was purified using AMPure XP (PCR product: AMPure XP beads = 1:0.8). The purified and 10-fold diluted 1st PCR products were used as templates for the 2nd PCR. The 2nd PCR was carried out with a 20-$\mu$l reaction volume containing 10 $\mu$l of 2 × Platinum SuperFi II PCR Master Mix, 4.0 $\mu$l of each 2.5 $\mu$M 1st-indexing-2nd-PCR F/R primer, and 2.0 $\mu$l of diluted 1st PCR product. The reaction volume for the 2nd PCR in “the 1st PCR indexing with Platinum” was increased so that the yield of 2nd PCR amplicons was sufficient (note that there was only one composite sample after the 1st PCR, unlike in the 2nd PCR indexing method).

The 3 libraries (i.e. one library for each treatment) were purified using AMPure XP (PCR product: AMPure XP beads = 1:0.8; Beckman Coulter, Brea, CA, USA). Target-sized DNA of the purified library (ca. 356 bp) was excised using E-Gel SizeSelect (ThermoFisher Scientific, Waltham, MA, USA). The double-stranded DNA concentrations of the libraries were quantified using a Quantus Fluorometer (Promega, Madison, WI, USA), and the 3 libraries were combined as one sample. The double-stranded DNA concentration of the combined library was then adjusted to 50 pM using 10 mM Tris-HCl (pH 8.5) and the DNA was then sequenced by the iSeq 100 system (Illumina, San Diego, CA, USA) using iSeq 100 Reagent v2 (2 × 150 bp PE). For the sequencing, 30% PhiX was spiked-in to improve the sequencing quality.

**Experiment II: Library preparation protocols and iSeq sequencing**

For 36 samples, the 1st PCR was performed by the 1st PCR indexing protocol using Platinum as described in library preparations in Experiment I except that one 1st PCR replicate with 4 $\mu$l of DNA template was employed (note that this modification is appropriate based on the results of Experiment III). For the “with exonuclease” treatment, 4 $\mu$l of ExoSAP-IT Express was added to each 10-$\mu$l PCR product. The mixture was incubated for 4 min at 37°C, followed by the incubation for 1 min at 80°C. For the “without exonuclease” treatment, the exonuclease purification was omitted. Then, the 3 indexed samples were combined as one composite sample which was incubated for 5, 30, or 120 min on ice or at room temperature. The 12 pooled libraries (= [with/without exonuclease] × [2 temperatures] × [3 durations]) were purified using AMPure XP as described in library preparations in Experiment I. The 2nd PCR was performed as described in library preparations in Experiment I except that “2nd-indexing-2nd-PCR primers” were used to append additional indices to the library (i.e., “quad-index approach”). This approach was adopted because the 3 indices used to distinguish the 3 samples were identical for 12 treatments.

The 12 libraries were purified, target-sized DNA of the purified library was excised, the double-stranded DNA concentrations of the libraries were quantified, and the 12 libraries were combined as in Experiment I. The double-stranded DNA concentration of the combined library was then adjusted to 50 pM using 10 mM Tris-HCl (pH 8.5) and the DNA was then sequenced by the iSeq 100 using iSeq 100 Reagent v2 (2 × 150 bp PE). For the sequencing, 30% PhiX was spiked-in to improve the sequencing quality.

**Experiment III: Library preparation protocols and iSeq sequencing**

DNA libraries were prepared by following the method described in library preparations in Experiment I with some modifications. For “replicate” treatment, the 1st PCR carried out with a 10-$\mu$l reaction volume containing 5.0 $\mu$l of 2 × Platinum SuperFi II PCR Master Mix, 2.0 $\mu$l of each 2.5 $\mu$M 1st-indexing-1st-PCR F/R primer (the final concentration was 0.5 $\mu$M), and

1.0 $\mu$l of template. The number of the 1st PCR replicates was 1, 2, 4, or 8, and the replicates were combined after the 1st PCR. Thermal cycle profiles were the same as in Experiment I and II. After the 1st PCR, 10 $\mu$l of the pooled library was purified by adding 4 $\mu$l of ExoSAP-IT Express as described in Experiment II.

For "volume" treatment, the first PCR was carried out with a 20-$\mu$l reaction volume to enable including 8 $\mu$l of template DNA. The reaction mixture contained 10.0 $\mu$l of 2 × Platinum SuperFi II PCR Master Mix, 1.0 $\mu$l of each 10 $\mu$M 1st-indexing-1st-PCR F/R primer (final concentration is 0.5 $\mu$M each), $H_2O$ and template DNA. The volume of the template DNA was 1 $\mu$l, 2 $\mu$l, 4 $\mu$l, or 8 $\mu$l, and the volume of $H_2O$ was adjusted so that the total volume of the reaction was 20 $\mu$l. Thermal cycle profiles were the same as in Experiment I and II. After the 1st PCR, 10 $\mu$l of the pooled library was purified by adding 4 $\mu$l of ExoSAP-IT Express as described in Experiment II.

The 12 1st PCR indices were used to distinguish the total 12 samples in 1-replicate and 1-$\mu$l treatments (2 sample types × [5 technical replicates + 1 negative control]). Similarly, the same 12 1st PCR indices were used in 2-replicate and 2-$\mu$l, 4-replicate and 4-$\mu$l, and 8-replicate and 8-$\mu$l treatments. Then, after the 1st PCR, 12 samples were combined as one sample, and we had eight libraries (each contained 12 samples) before the 2nd PCR. The 2nd PCR was performed as described in library preparations in Experiment I except that "2nd-indexing-2nd-PCR primers" were used to append additional indices to the library (i.e., quad-index approach). This approach was adopted to distinguish the 4 libraries.

After the 2nd PCR, the eight libraries were purified, target-sized DNA of the purified library was excised, the double-stranded DNA concentrations of the libraries were quantified, and the eight libraries were combined as one sample. The volume of each library combined was adjusted to normalize the concentrations of each 2nd PCR product. The double-stranded DNA concentration of the combined library was then adjusted to 50 pM using 10 mM Tris-HCl (pH 8.5) and the DNA was then sequenced by the iSeq 100 system using iSeq 100 Reagent v2 (2 × 150 bp PE). For the sequencing, 25% PhiX was spiked-in to improve the sequencing quality.

### Sequence data processing

As described in the main text, our sequence data includes sample-specific indices inside the sequencing primers and we prepared a custom shell script based on seqkit (Shen *et al.* 2016) to demultiplex our samples (https://github.com/ong8181/eDNA-early-pooling; see also https://doi.org/10.5281/zenodo.6045851). Raw sequences were demultiplexed and MiFish primer regions were trimmed by cutadapt (Martin 2011). The quality of our sequence data was high, and after primer trimming, 8,948,113 sequence reads (%>Q30 = 93.9; average 43,863 reads per sample) remained for the three experiments.

The demultiplexed, primer-trimmed sequences were processed using `DADA2` (Callahan *et al.* 2016), an amplicon sequence variant (ASV) approach, for each iSeq run. At the quality filtering process, low quality and unexpectedly short reads were removed using `DADA2::filterAndTrim()` function with arguments of `truncLen = c(110,110)`, `minLen = 100`, and `maxEE = c(2,2)`. Error rates were learned using `DADA2::learnErrors()` function. Then, sequences were dereplicated, error-corrected using `DADA2::dada()` with an argument `pool = TRUE`, and merged to produce an ASV-sample matrix. Chimeric sequences were removed using the `DADA2::removeBimeraDenovo()` function. Then, ASVs detected in the three experiments were merged and clustered into OTU at 97% similarity using `DECIPHER` package of R (Wright 2016), which converted the ASV-sample matrix into the OTU-sample matrix.

Taxonomic identification was performed for OTUs based on the query-centric auto-*k*-nearest-neighbor (QCauto) method (Tanabe & Toju 2013) and subsequent taxonomic assignment with the lowest common ancestor algorithm (Huson *et al.* 2007) using "overall" database and `clidentseq` and `classigntax` commands implemented in `Claident` (https://www.claident.org/). Because the QCauto method requires at least two sequences from a single microbial taxon, standard DNAs were separately identified using BLAST (Camacho *et al.* 2009). Scripts to analyze the sequence data are available at https://github.com/ong8181/eDNA-early-pooling.
