## Supplementary Tables and Figures for "An efficient early-pooling protocol for environmental DNA metabarcoding"

### Contents

- Table S1. Standard DNA sequences.
- Table S2. Thermal cycle profiles for PCR in Experiment I.
- Table S3. Cost and time estimations for the common 2nd PCR indexing protocol and the early-pooling protocol.
- Figure S1. Raw sequence reads generated in Experiment 1 (not rarefied).
- Figure S2. Relative abundance and the number of OTUs detected in Experiment I.
- Figure S3. Effects of the library preparation methods on the detected relative abundance of the three most-dominant OTUs.
- Figure S4. Relative abundance of sequence reads detected in Experiment III.
- Figure S5. Effects of library preparation protocols on the community composition of fish eDNA detected in the Nagahama samples.

**Table S1. Fish-like standard DNA sequences**

| Sequence name | Sequence |
|---|---|
| STD_MiFish02 | GTCGGTAAAACTCGTGCCAGCCACCGCGGTTATACGACAGGCCCAAGTTGATCTTG<br>AACGGCGTAAAGAGTGGTTAGATTTCCCTACTGCTAAAGCCGAAGCACCGCCGTGC<br>TGTTATACGTGCCTCACAGTGAGAAGGGCGAAAACGAAAGTAGCTTTATTTCCGTC<br>ACGCGAACCCACGAAAGCTAAGAAACAAACTGGGATTAGATACCCCACTATG |
| STD_MiFish04 | GTCGGTAAAACTCGTGCCAGCCACCGCGGTTATACGACAGGCCCAAGTTGATATCC<br>CACGGCGTAAAGAGTGGTTAGAACCGGAAACGCGTAAAGCCGAAGAACATCAGTGC<br>TGTTATACGCATTCGATTAGGTGAATTTCAGTAACGAAAGTAGCTTTACCGATCAA<br>ATCCGAACCCACGAAAGCTAAGAAACAAACTGGGATTAGATACCCCACTATG |
| STD_MiFish05 | GTCGGTAAAACTCGTGCCAGCCACCGCGGTTATACGACAGGCCCAAGTTGACTGGT<br>GTCGGCGTAAAGAGTGGTTAACTAACGATCATTATAAAGCCGAAATTCCCCTAAGC<br>TGTTATACGTCAAATCAAAAAGGAATCGCTGGTACGAAAGTAGCTTTAGTTATGAT<br>GCATGAACCCACGAAAGCTAAGAAACAAACTGGGATTAGATACCCCACTATG |
| STD_MiFish08 | GTCGGTAAAACTCGTGCCAGCCACCGCGGTTATACGACAGGCCCAAGTTGACTTAG<br>CCCGGCGTAAAGAGTGGTTATTTGAATTAAGATCTAAAGCCGAATTTCGACGCAGC<br>TGTTATACGGGTTCTGGATTCAGAATGCCCGATACGAAAGTAGCTTTACCAGTGCG<br>ATTAGAACCCACGAAAGCTAAGAAACAAACTGGGATTAGATACCCCACTATG |
| STD_MiFish09 | GTCGGTAAAACTCGTGCCAGCCACCGCGGTTATACGACAGGCCCAAGTTGACGTAC<br>TACGGCGTAAAGAGTGGTTACGTGTCGCACCTGGTAAAGCCGAAGCCCCTCTAGGC<br>TGTTATACGAGACAGAATCGGCGAAGCTCGCGCACGAAAGTAGCTTTAGCGGAGAT<br>ATGCGAACCCACGAAAGCTAAGAAACAAACTGGGATTAGATACCCCACTATG |
| STD_lateolabrax | GCCGGTAAAACTCGTGCCAGCCACCGCGGTTATACGAGGGGCCCAAGTTGAATATA<br>AACGGCGTAAAGGGTGGTTAAGCCACACCCTACTGGTAAAGCCGAATCTCGCTCAC<br>GCTGTTATACGTGAGCCATATCGAGAAAGACTCCAACGAAAGTGGCTTTACCACAT<br>TTGAACCTACGAAAGCTGGGGCACAAACTGGGATTAGATACCCCACTATG |
| STD_takifugu | GTCGGTAAAACTCGTGCCAGCCACCGCGGTTATACGAGAGACCCAAGTTGAATTGA<br>GTCGGCGTAAAGGGTGGTTAACGCGTCTAGCGTCATGATGAGACCGAATGTCCTCC<br>TCGCTGTTATACGTGTAATGATTTCCGAACCTCTTAAACGAAAGTAGCCTCACTAA<br>CTCGAACCCACGAAAGCTAGGACACAAACTGGGATTAGATACCCCACTATG |
| STD_cyprinus | GCCGGTAAAACTCGTGCCAGCCACCGCGGTTAGACGACAGGCCCTAGTTGAGCCAC<br>ACCCGGCGTAAAGGGTGGTTAGGACGGGCGACGTTTAAAGTCAAAGAAGCTCCATG<br>CCGTTATACGATGGTTAGTTCCGGAACGGCAGGAACGAAAGTAACTTTATAAGTGT<br>ATGCCGAACCCACGAAAGCTGAGAAACAAACTGGGATTAGATACCCCACTATG |
| STD_Engraulis | GCCGGTAAAACTCGTGCCAGCCACCGCGGTTATACGAGAGACCCTAGTTGAGGGTT<br>TCGGCGTAAAGAGTGGTTATGCTGGCGTATCATTTAAAGCAGAATGAGTGCGCCAC<br>TGTTATACGGCGCCGATAAATCGAATACCGTTGACGAAAGTGACTTTACAGGGGCC<br>ACCTGGAAGCCACGAAAGCTGGGACACAAACTGGGATTAGATACCCCACTATG |
| STD_Anguilla | GCCGGTAAAACTCGTGCCAGCCACCGCGGTTATACGAGGGGCTCAAATTGAGTTGT<br>GGCGGCGTAAAGCGTGATTATTGACCTCATCTCGTAAAGCCAAATTTCCGCGTAGC<br>TGTCATACGAAGTACCGGCTGGGGAAACTCAGGGACGAAAGTGGCTTTAGGTAGGA<br>GGAACCCACGACAGTTGAGAAACAAACTGGGATTAGATACCCCACTATG |

**Table S2. Thermal cycle profiles for PCR in Experiment I**

| PCR step | 2nd PCR indexing with KAPA | 2nd PCR indexing with Platinum | 1st PCR indexing with Platinum |
|---|---|---|---|
| **1st PCR** | | | |
| Initial denaturation | 98°C, 3 min | 98°C, 30 sec | 98°C, 30 sec |
| Denature | 98°C, 20 sec | 98°C, 10 sec | 98°C, 10 sec |
| Annealing | 65°C, 15 sec | 60°C, 10 sec | 60°C, 10 sec |
| Extension | 72°C, 15 sec | 72°C, 15 sec | 72°C, 15 sec |
| (No. of Cycle) | (× 35) | (× 35) | (× 35) |
| Final extension | 72°C, 5 min | 72°C, 5 min | 72°C, 5 min |
| Cooling | 4°C | 4°C | 4°C |
| **2nd PCR** | | | |
| Initial denaturation | 98°C, 3 min | 98°C, 30 sec | 98°C, 30 sec |
| Denature | 98°C, 20 sec | 98°C, 10 sec | 98°C, 10 sec |
| Annealing + Extension | 72°C, 15 sec | 72°C, 15 sec | 72°C, 15 sec |
| (No. of Cycle) | (× 12) | (× 12) | (× 12) |
| Final extension | 72°C, 5 min | 72°C, 5 min | 72°C, 5 min |
| Cooling | 4°C | 4°C | 4°C |

**Table S3. Cost and time estimations for the common 2nd PCR indexing protocol and the early-pooling protocol**

| Step | Assumptions | Price/Time for 96 samples[1] |
|---|---|---|
| **2nd-PCR-indexing** | | |
| 1st PCR | 10-$\mu$l scale, 4 replicates per sample, KAPA HiFi HotStart ReadyMix used | \$180 (~ 3 hours) |
| Purification | AMPure XP, 16 $\mu$l for each sample | \$130 (~ 1 hours) |
| 2nd PCR | 20-$\mu$l scale, 1 replicate per sample, KAPA HiFi HotStart ReadyMix used | \$89 (~ 2 hours) |
| Pooling, Purification | AMPure XP, 16 $\mu$l for 1 sample | \$1 (~ 30 min) |
| **Total** | | **\$ 400 per 96 samples (up to 6 hours 30 min)** |
| **1st-PCR-indexing** | | |
| 1st PCR | 20-$\mu$l scale, 1 replicate per sample, Platinum SuperFi II DNA polymerase used | \$51 (~ 2 hours) |
| Purification, Pooling | Exonuclease, 2 $\mu$l for each sample | \$91 (~ 30 min) |
| Adjust concentration | AMPure XP, 16 $\mu$l for 1 sample | \$1 (~ 30 min) |
| 2nd PCR | 20-$\mu$l scale, 1 sample per library, Platinum SuperFi II DNA polymerase used | \$1 (~ 1 hours) |
| Purification | AMPure XP, 16 $\mu$l for 1 sample | \$1 (~ 30 min) |
| **Total** | | **\$ 144 per 96 samples (up to 4 hours 30 min)** |

[1]Cost and time estimations are approximate and may change depending on the detailed experimental protocols. In addition, costs for plastic consumables are not included.

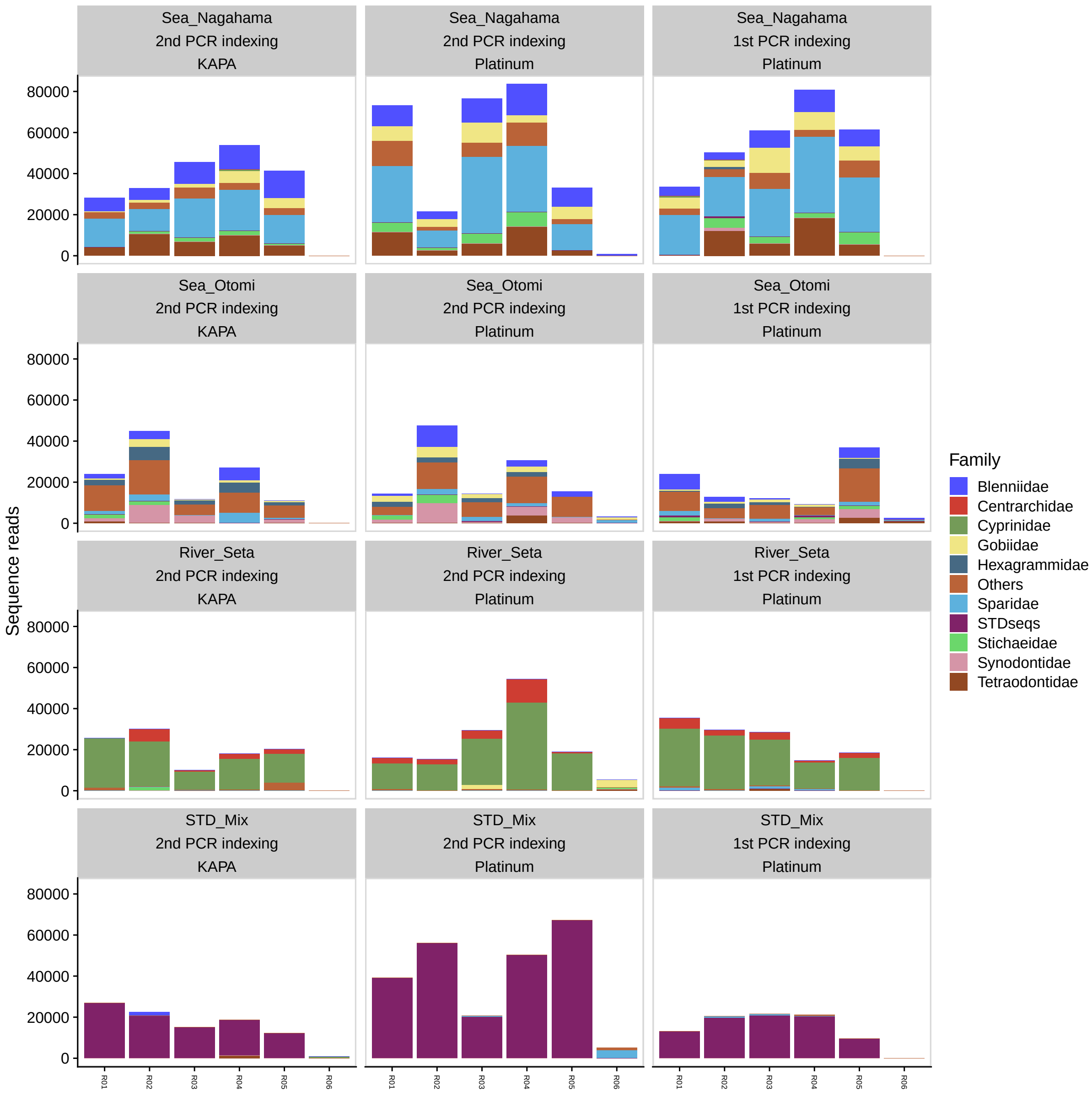

**Figure S1. Sequence reads generated in Experiment I (not normalized)**. Each panel shows sequence reads for 5 positive samples (R01-R05) and 1 negative sample (R06; $H_2O$). Colors indicate fish family assigned by `Claident`, or the standard fish DNAs.

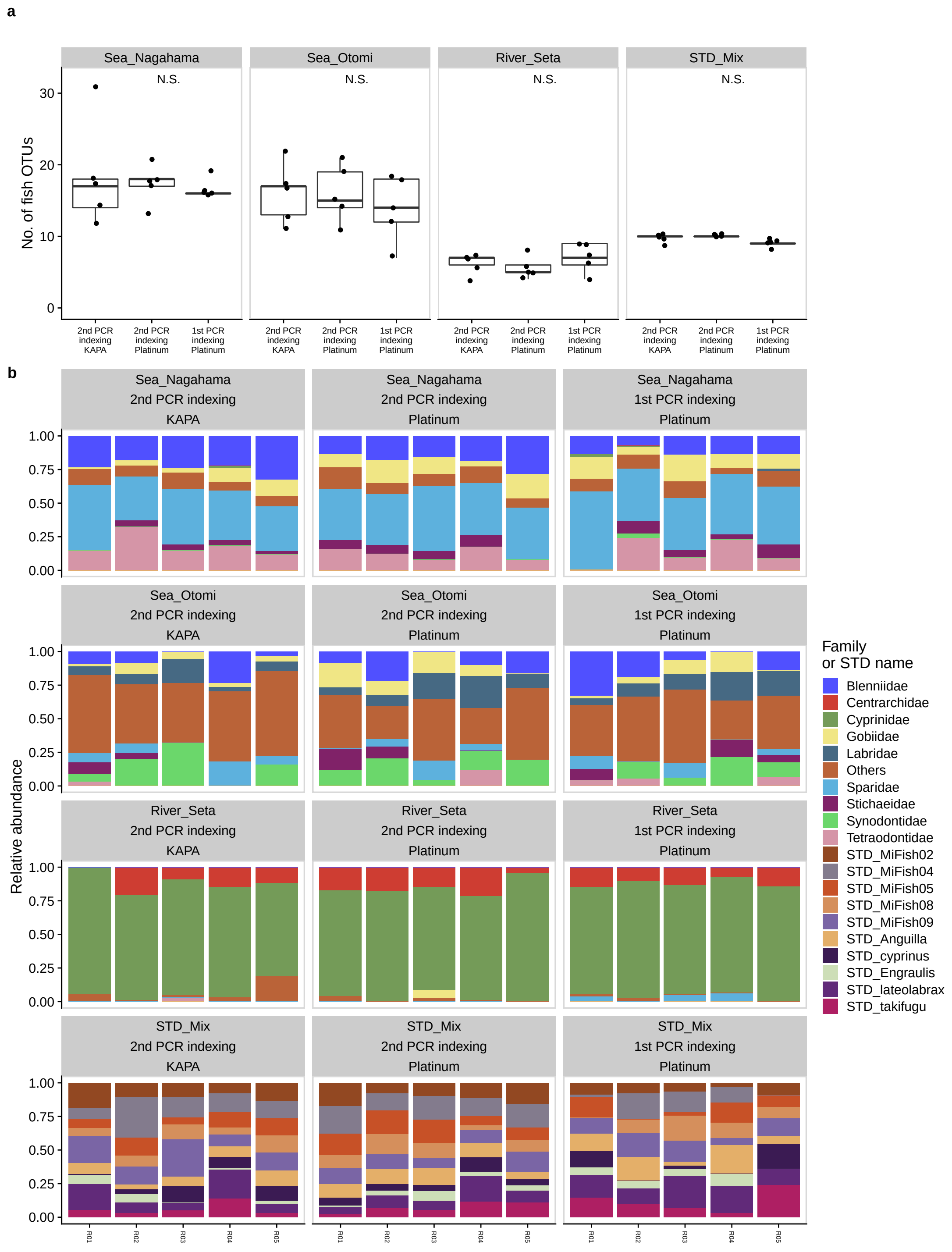

**Figure S2. Relative abundance and the number of OTUs detected in Experiment I.** (**a**) The number of OTUs detected in each treatment. *x*-axis indicates experimental protocols (i.e., the 1st or 2nd PCR indexing protocol and KAPA HiFi HostStart ReadyMix or Platinum SuperFi II PCR Master Mix). (**b**) Relative abundance of OTUs detected in each treatment. *x*-axis indicates replicates in the treatment. Colors indicate fish family assigned by `Claident`, or the standard fish DNAs.

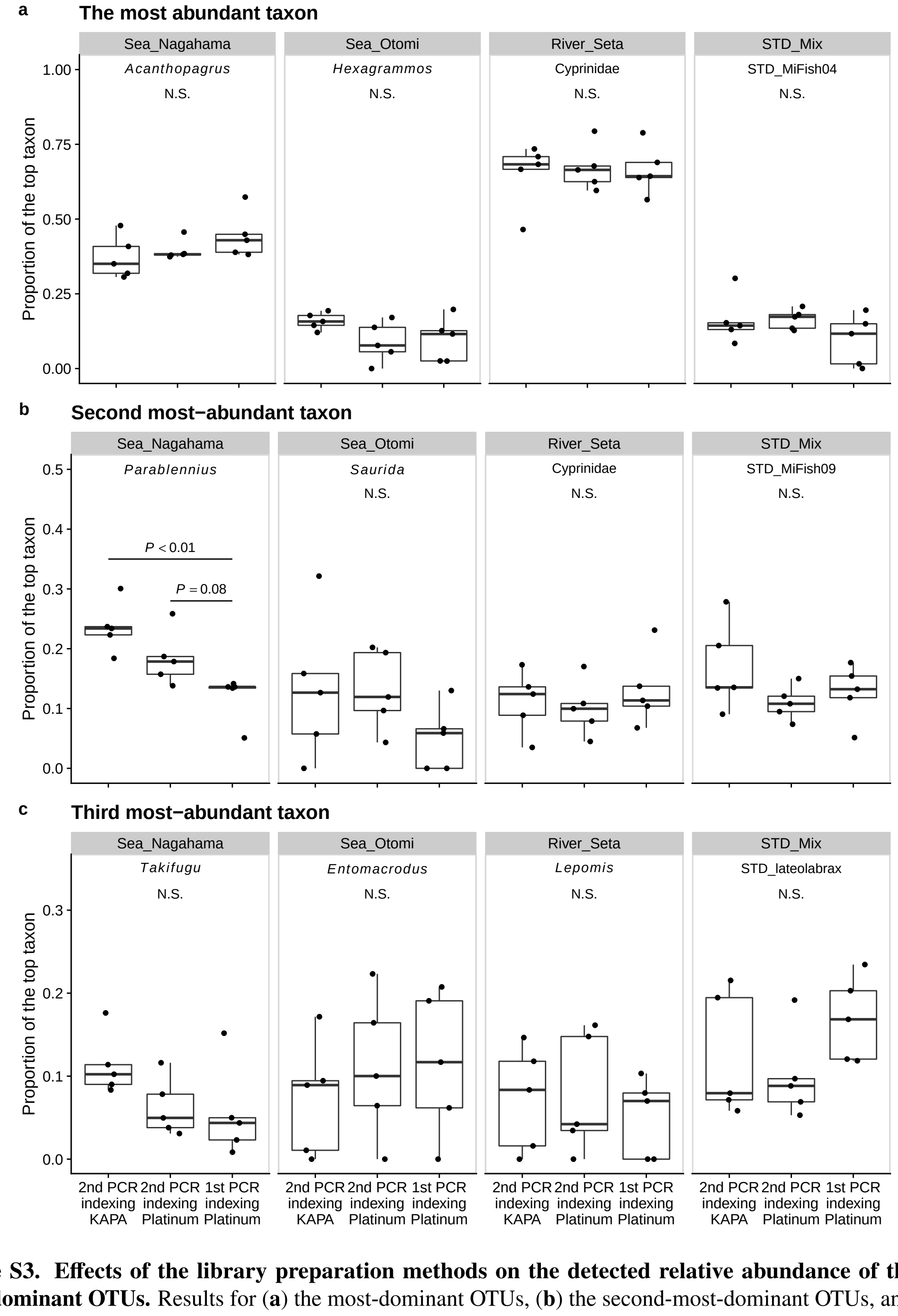

**Figure S3. Effects of the library preparation methods on the detected relative abundance of the three most-dominant OTUs.** Results for (**a**) the most-dominant OTUs, (**b**) the second-most-dominant OTUs, and (**c**) the third-most-dominant OTUs. Note that the taxon that each OTU represents is different depending on the study site. Taxon name assigned to each OTU is shown at the top of each panel. Statistical clarity was tested by GLM that assumed normal distributions of the sequence reads (see Methods).

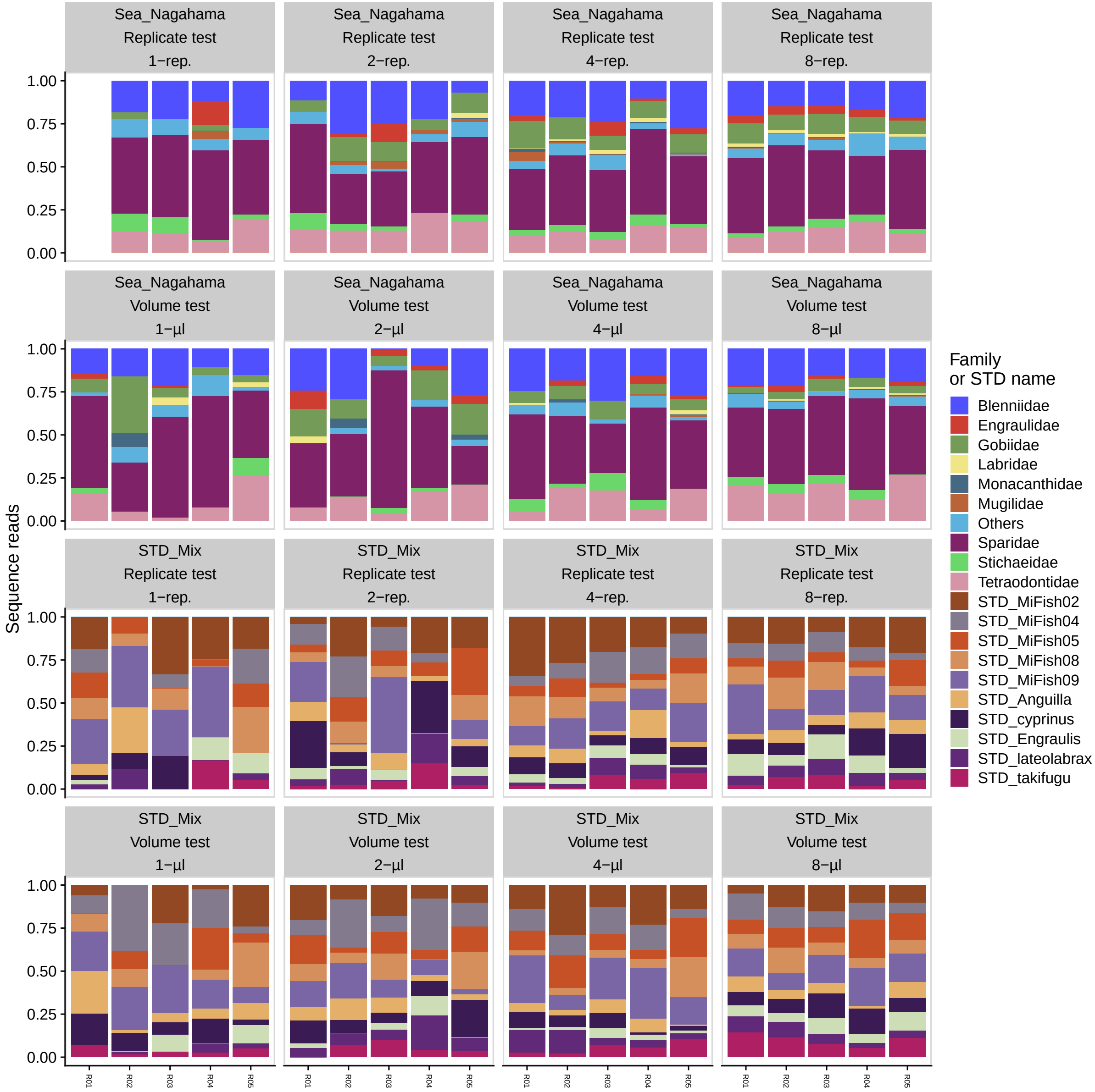

**Figure S4. Relative abundance of sequence reads detected in Experiment III.** Each panel indicates 5 replicates for each treatment. Note that R01 in 1-rep. treatment in the replication test was removed because only 304 reads were assigned to the sample. Colors indicate fish family assigned by `Claident`, or the standard fish DNAs.

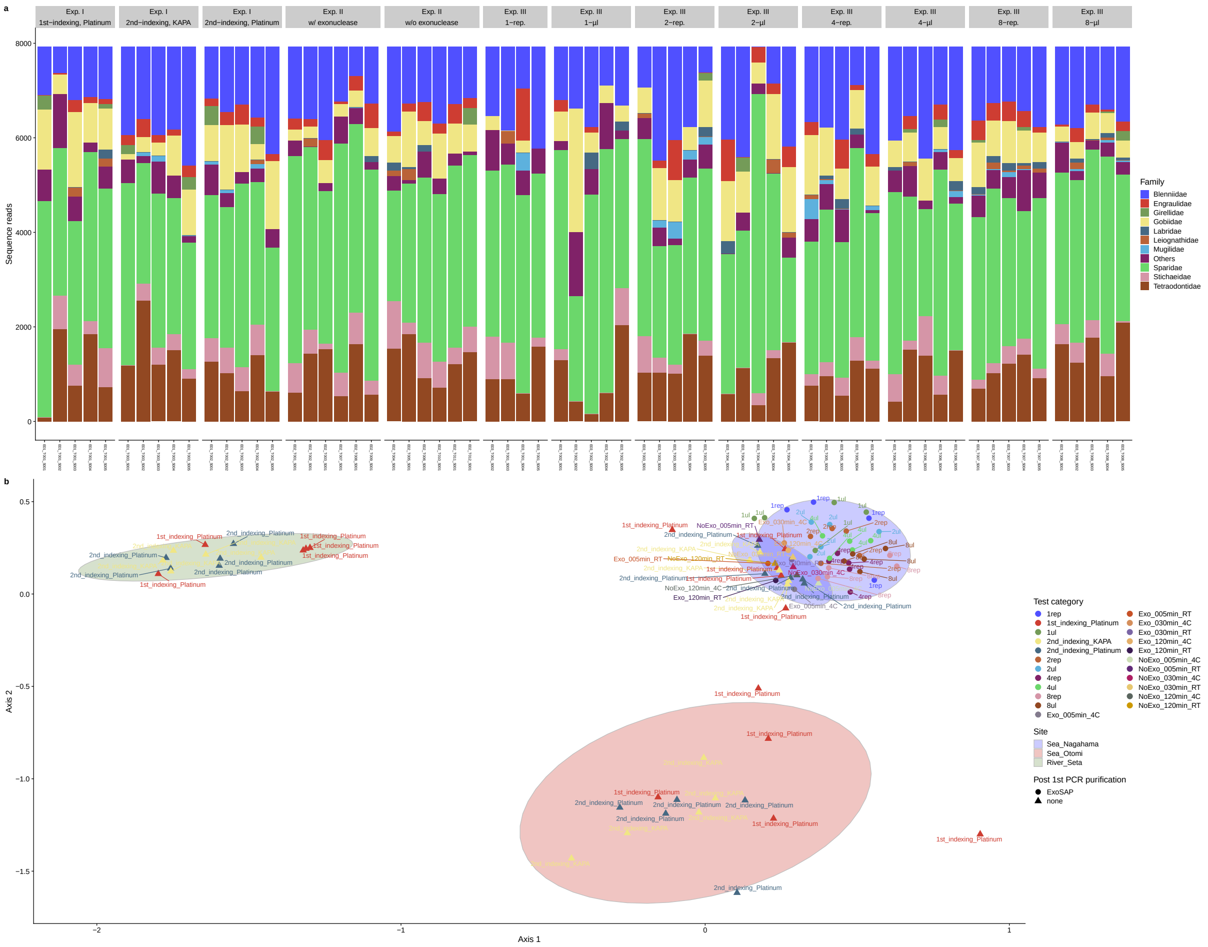

**Figure S5. Effects of library preparation protocols on the community composition of fish eDNA detected in the Nagahama samples.** (**a**) Relative abundance of sequence reads detected in the Nagahama samples. Each panel shows the indicated experimental treatment, and colors indicate fish family. (**b**) Nonmetric dimensional scaling (NMDS) of all natural eDNA samples analyzed in the present study. Various library preparation methods were tested for the Nagahama samples, and all Nagahama samples were clearly distinguished from natural eDNA samples from the other study sites. Symbols and colors indicate the purification protocol after the 1st PCR and experimental treatments, respectively. Ellipses indicate 95% confidential intervals for each study site and purification protocol.
